# Atomic structure and dynamics of the mechanosensitive channel MscL from *E. coli* by cryo-EM and solid-state NMR

**DOI:** 10.64898/2026.04.14.718163

**Authors:** Taoran Xiao, Alexandra Kovinko, Chaowei Shi, Henry Sawczyc, Denis Qoraj, Carl Öster, Thiemo Sprink, Sascha Lange, Spyridon Kosteletos, Han Sun, Daniel Roderer, Shanshuang Chen, Adam Lange

## Abstract

Mechanosensitive channels are central to cellular responses to membrane tension, yet the structural basis of their gating remains incompletely understood. Here, we determine the structures of wild-type and G22S mutants of MscL from *E. coli* (*Ec*MscL) by cryo-EM in peptide-based lipid nanodiscs and complement them with solid-state NMR measurements in liposomes to capture their dynamics in a native-like membrane environment. The cryo-EM structures reveal a closed conformation, whereas analysis of the low-threshold G22S mutant by NMR uncovers widespread conformational changes in both cytoplasmic and periplasmic regions. These data indicate enhanced dynamics and conformational heterogeneity in the mutant, revealing the early transitions from the closed towards the open state. Together, our results establish a synergistic framework integrating cryo-EM and NMR to resolve both structure and dynamics of mechanosensitive channels, and identify lipid-protein interactions as key determinants of MscL gating and mechanosensitivity. Our study further provides a quantitative benchmark for computational investigations of mechanogating and lays the foundation for the rational design of channels with tunable gating kinetics.

**Teaser:** By integrating cryo-EM and solid-state NMR, we reveal how lipid-coupled dynamics prime MscL for opening, capturing the earliest transitions from closed to active states.

## Introduction

Mechanosensitive (MS) channels are transmembrane proteins universally present in both prokaryotic and eukaryotic organisms (*1*), where they serve as fundamental components of cellular mechanotransduction systems. These channels function as direct sensors of mechanical forces, converting external and internal physical stimuli into biochemical signals that regulate crucial physiological processes. Through this function, MS channels enable cells to maintain structural integrity and adapt to rapidly changing environmental or intracellular conditions. In prokaryotic systems, MS channels play a pivotal role in osmoregulation. The opening of MS channels under hypo-osmotic shock facilitates the rapid release of cytoplasmic solutes, serving as a safety valve to prevent cellular rupture and death (Fig. 1A I-III) (*2*–*5*). In eukaryotic cells, mechanosensitive channels regulate key physiological and developmental processes such as the sensation of touch, hearing, pain, bone and muscle homeostasis regulation, and form part of complex mechanotransductive networks that coordinate cell communication and tissue organization (*6*–*9*).

**Fig 1.**
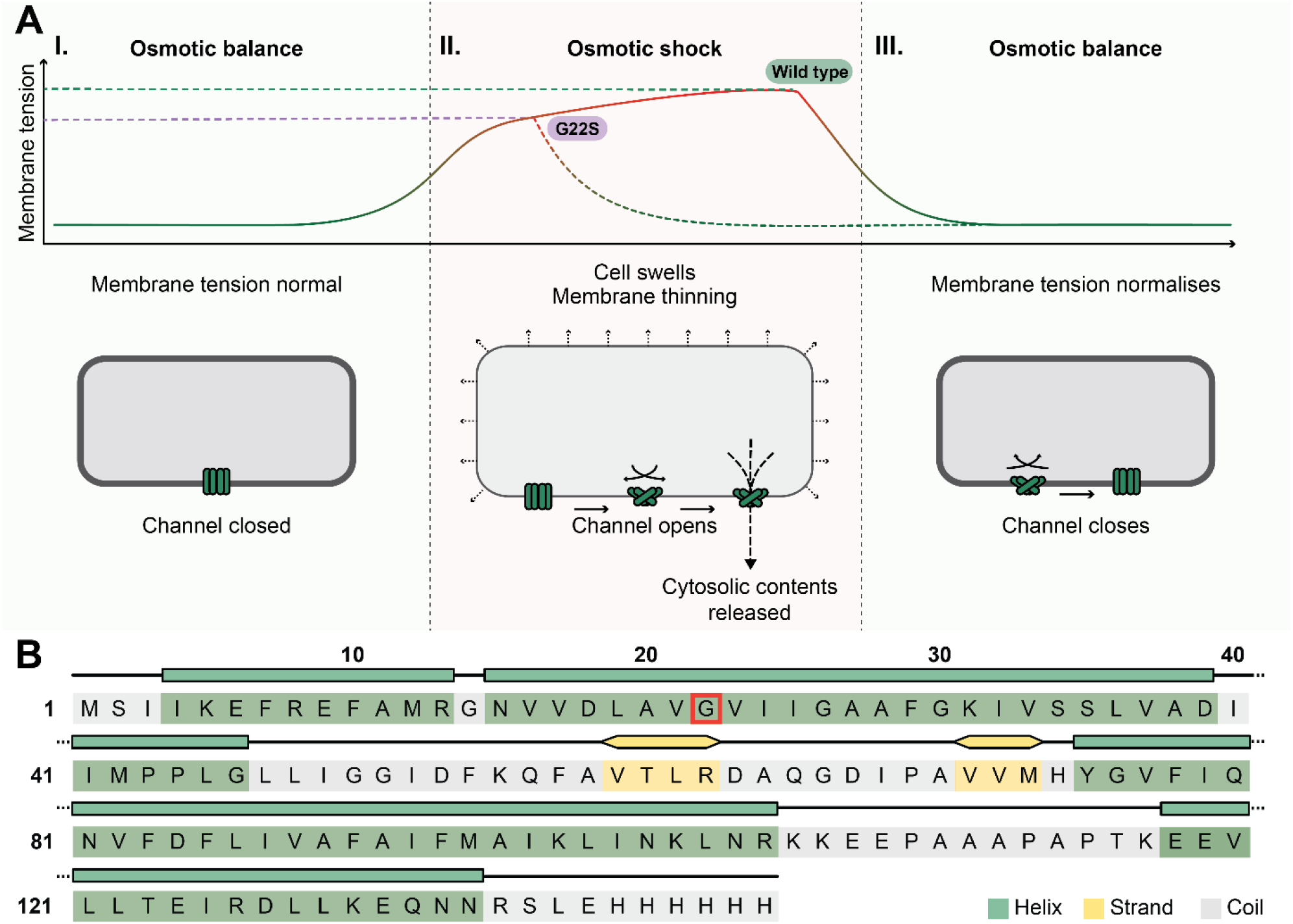
Biological function and secondary structure prediction of the mechanosensitive channel of large conductance in *E. coli*. (*Ec*MscL) (**A**) Schematic diagram of the opening of *Ec*MscL as membrane tension increases, with the lower threshold of tension required to open the G22S mutant compared to wild type (WT) **I**.) *Ec*MscL channels are embedded in the inner membrane of *E. coli*, under normal membrane tension the channels are closed **II**.) An increase in membrane tension opens *Ec*MscL during osmotic shock, releasing the cytosolic contents in a non-selective way to avoid cell lysis. **III**.) The channel closes again when the osmotic balance is recovered. (**B**) Amino acid sequence and secondary structure prediction of *Ec*MscL according to *Mt*MscL by PSIPRED 4.0 (*21*) (green, α-helical; yellow, β-sheet; grey, random coil/unstructured; red rectangle, mutation site in G22S mutant).

Due to their conserved and essential function, MS channels have been the focus of many studies. The focus of this work is primarily on prokaryotic MS channels, for an overview of the research on eukaryotic MS channels and their function, please see refs (*10, 11*). In terms of prokaryotes, the mechanosensitive channel of large conductance (MscL) from *Mycobacterium tuberculosis* (*Mt*MscL) was the first MS channel to be structurally characterized by X-ray crystallography nearly three decades ago (*12*). The structure of *Mt*MscL revealed the channel as a homopentamer, with each monomer containing a helical N-terminal domain (NTD), two transmembrane domains (TM1 and TM2), and a helix in the C-terminal domain (CTD). The periplasmic loop (PL) connects TM1 and TM2, and a further loop (L) connects TM2 and the CTD (fig. S1). The *Mt*MscL structure, and others that have followed it, have since been widely used as a model system to elucidate the mechanism of mechanosensation through numerous studies using electrophysiology (*13, 14*), biochemistry (*15*), genetics (*16*), and molecular dynamics simulations (*3, 17*). The initial *Mt*MscL structure (solved in detergent) has been essential for understanding the structural basis of bacterial MS channels. However, structural information in the presence of a lipid bilayer is essential for the elucidation of the central lipid-based mechanotransduction mechanism (*18*). This is especially poignant as MS channels appear to not only sense tension of the membrane as a whole, but also more subtle interactions between the protein and membrane lipids (*19, 20*).

Solid-state NMR has become a powerful approach in the last decade for investigating membrane proteins in native-like environments, enabling the characterization of conformations directly in complex lipid bilayers (*22, 23*). Recent studies have demonstrated its ability to probe lipid–protein interactions and conformational equilibria at high resolution using paramagnetic and multinuclear techniques (*24, 25*), as well as to elucidate functional mechanisms in membrane systems, including transmembrane signaling and conformational plasticity (*26, 27*). Applications span a wide range of biological systems, from bacterial membrane complexes to viral ion channels, accessory proteins, and intramembrane enzymes (*28*–*31*), as well as viral maturation processes (*32*) and amyloid fibrils with varying toxicity and seeding potential (*33*), highlighting the versatility of the method. Furthermore, ssNMR enables the investigation of membrane protein stability, unfolding pathways, and functional switching (*34, 35*), while recent developments in dynamic nuclear polarization with fast magic angle spinning have significantly expanded the accessible sensitivity and timescales of these measurements (*36*).

Due to the intrinsic membrane interactions required for MS channel function, solid-state NMR (ssNMR) techniques have been used to explore the structure, function, and conformational dynamics of this class of channels in native-like liposomes.

Initial efforts of studying MscL using ssNMR employed cell-free expression coupled with selective isotope labeling to obtain high-quality spectra of the full-length *Ec*MscL channel embedded in lipid assemblies (*37*). This approach validated efficient strategies for producing and probing MscL in its native-like environment and confirmed that the protein produced by cell-free methods retains a fold consistent with native states. More recent studies were performed on MscL from *Methanosarcina acetivorans* (*Ma*MscL) reconstituted into liposomes (*38*) as well as in *E. coli* native membranes (*39*–*41*). Using these set-ups, extensive resonance assignments and detailed secondary structure information could be obtained for *Ma*MscL within lipid membranes. These studies revealed that the transmembrane architecture of MscL is largely conserved between reconstituted and native contexts, while also uncovering subtle differences at the interface with surrounding lipids. Complementary investigations integrating molecular dynamics simulations used ssNMR to map the interaction of small-molecule ligands such as the antibiotic Brilliant Green with MscL. These studies found that small molecules primarily interacted with the cytoplasmic entrance and gating residues, demonstrating ligand-induced shifts in conformational equilibria (*42*).

Despite the relative abundance of previous ssNMR studies and the good sequence homology of *Ec*MscL with MscL variants for which structures have been resolved (Fig. 1B and fig. S2), a complete structure of MscL from *E. coli* has remained elusive. Additionally, the relatively small size and intrinsic conformational flexibility have further limited high-resolution characterization of *Ec*MscL by both X-ray crystallography and cryogenic electron microscopy (cryo-EM). Here, leveraging recent advancements in membrane models, most notably peptide-based lipid nanodiscs, we present the structure of both wild type and G22S mutant of *Ec*MscL in nanodiscs (MscL-ND) by single particle cryo-EM to a high resolution. Complementing these high-resolution structures, ssNMR measurements of both WT and G22S mutant of *Ec*MscL in native-like liposomes, enabled the assignment and subsequent structural analysis of a total of 57 out of 136 residues primarily spanning the periplasmic loop, the CTD, as well as several transmembrane residues. In carbon-detected spectra of protonated wild-type channel, assignments are more sparse for TM2 compared to TM1, pointing to structural heterogeneity of the lipid-facing TM2. Whilst the cryo-EM structures show no major global differences between the WT and G22S mutant structures, NMR uncovers pronounced, region-specific changes in conformational dynamics, with enhanced flexibility of the PL seen for the mutant as well as a slight stabilization of the CTD. Together, these results highlight the power of integrating cryo-EM and ssNMR to elucidate not only the static structures of membrane proteins in native-like environments, but also the lipid-coupled conformational dynamics that underpin their structure-function relationship.

## Results

### Overall cryo-EM structures of wild-type *E. coli* MscL and the G22S mutant

The 3D structures of full-length wild-type (WT) and G22S mutant of *Ec*MscL channel reconstituted in nanodiscs were determined using single-particle cryo-EM, achieving a global resolution of 3.7 Å and 3.5 Å, respectively (Fig. 2). Though buried in the lipid bilayer, the channel core, particularly the transmembrane helices, is clearly visible in the cryo-EM density map. Reconstructions with C5 symmetry resulted in similar maps with those reconstructed without applied symmetry (C1) besides improved map quality and overall resolution, suggesting that *Ec*MscL is indeed 5-fold related (Fig. 2C). Because *de novo* model building is not feasible due to limited resolution, structural model predicted by Alphafold3 (*43*) (fig. S1) were fitted into the electron density map, and registration was validated by strong density for aromatic side chains. The initial models were refined against the reconstructions to achieve good model geometry. Notably, despite that the Alphafold3 prediction of the G22S mutant indicates an expanded form (fig. S1), we chose the WT structure as the starting model because it better agrees with the experimental EM map of the G22S mutant, and subsequently refined it similarly as for WT. Whereas the overall topology and architecture of *Ec*MscL resemble those of the homologous *Mt*MscL determined by X-ray crystallography in detergent, substantial differences are observed, particularly in the periplasmic region, the cytoplasmic end of TM2, and the cytoplasmic region (fig.S1). It is possible that these deviations represent evolutionary divergence between homologs, although influences from structural determination conditions such as the membrane mimetic environment can not be ruled out yet.

**Figure 2.**
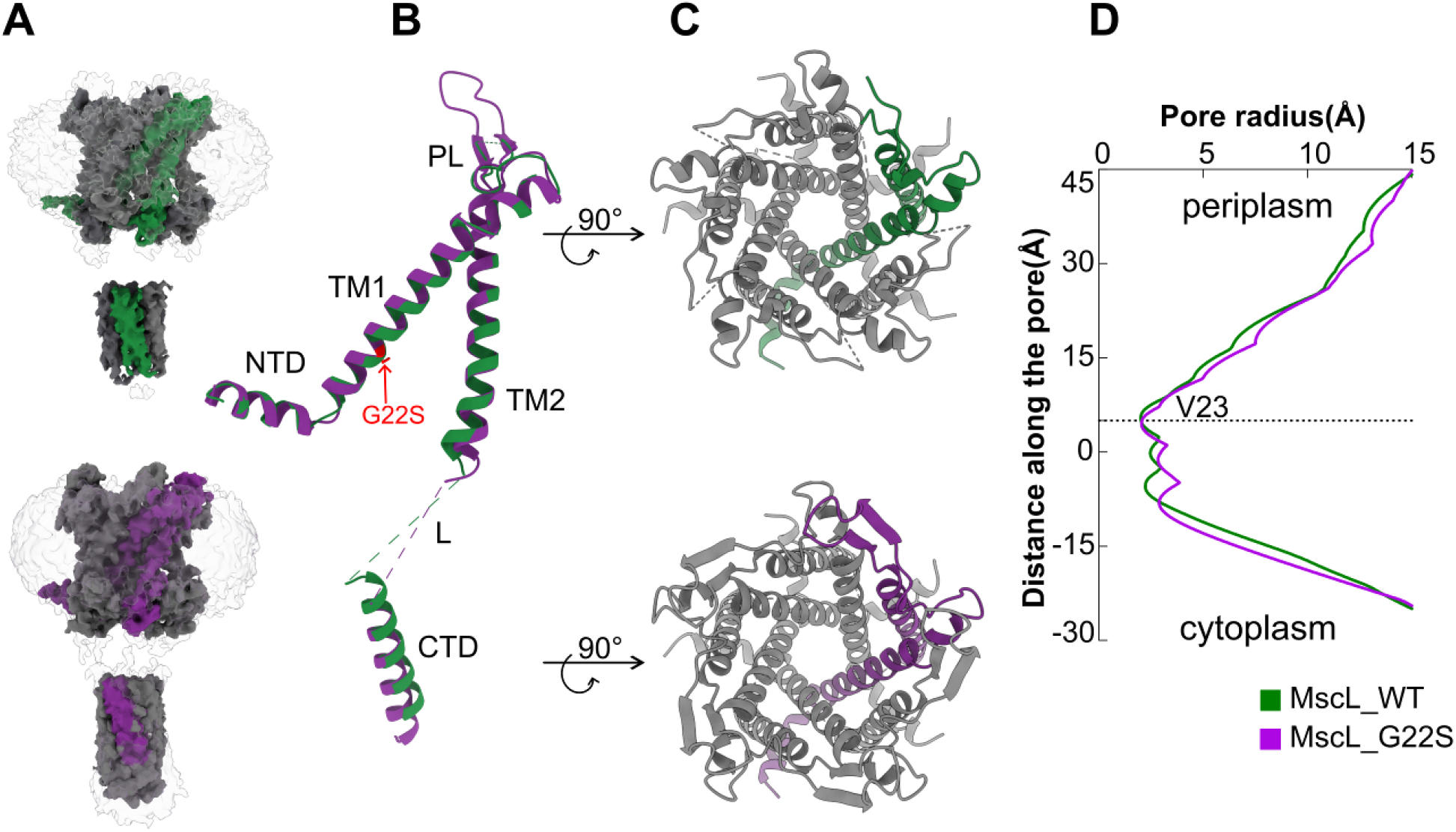
The overall cryo-EM structures of wild-type *Ec*MscL and its G22S mutant. (**A**) Cryo-EM reconstruction of the wild-type (top) and G22S mutant (bottom) full-length channel in nanodiscs (side-on) with a single subunit uniquely colored (green: WT, purple: G22S). (**B**) An overlay of monomeric subunits of the wild-type and G22S structures. NTD: N-terminal domain, TM1: transmembrane helix 1, PL: periplasmic loop, TM2: transmembrane helix 2, L: loop, CTD: C-terminal domain. The mutation site is highlighted in red. (**C**) Periplasmic (top-down) view of the cryo-EM structures of the wild-type (top) and G22S mutant (bottom) full-length channel, with the CTD hidden for clarity.(**D**) Comparison of the ion pore profiles of the wild-type (green) and G22S mutant (purple) channels, with the narrowest part, at V23, labeled.

The overall structures of wild-type *Ec*MscL and the G22S mutant are similar, with an all-atom root mean square deviation (RMSD) of 2.4 Å. Each *Ec*MscL protomer consists of an N-terminal domain (NTD) located on the cytoplasmic side of the membrane, two transmembrane helices (TM1 and TM2) connected by a large periplasmic loop (PL), and a C-terminal domain (CTD) composed of a remote helix connected to TM2 by a cytoplasimc loop (L) (Fig. 2B).

For the NTD, most of the main chain backbone could be fitted into the density and some bulky sidechains further confirmed the placement of the backbone (fig. S3). Interestingly, the NTD forms a short amphipathic helix containing 13 amino acids (1-13 MSIIKEFREFAMR), followed by a sharp turn mediated by G14 between the amphipathic helix and TM1. The pentameric arrangement of the transmembrane domain generates an hour-glass shaped channel pore that is broad on both the periplasmic and cytosolic periphery and is narrow near the center of TM1, where a hydrophobic restriction is formed (fig. S4). The pore cavity is primarily lined by the first transmembrane helices (TM1) and the peripheral TM2 interacts with the surrounding lipids (Fig. 2A).

The periplasmic loop (“PL”) is poorly resolved in contrast to the well-defined transmembrane regions, suggesting its flexibility, which has been implicated in modulating gating kinetics and mechanosensitivity (*44, 45*). However, the structural basis underlying its regulatory function is not well understood. The cryo-EM map supports the presence of a ω-loop and a short β-hairpin in the PL, consistent with a previous study (Fig. 4A) (*46*). However, the loop is unlikely to directly transmit mechanical force between the TM helices during channel activation because of its pronounced flexibility. Instead, accumulating evidence suggests a regulatory role in tuning gating sensitivity (*47*). Consistent with this notion, we complemented our cryo-EM investigation of WT *Ec*MscL with ssNMR experiments showing elevated hydrogen-deuterium exchange, which will be discussed later. How these dynamic properties translate into functional regulation of channel gating, however, remains an open question requiring further investigation.

The cytoplasmic region (residues 106-136) containing the cytoplasmic loop (L) and the CTD helical bundle is also less well resolved in both WT and G22S mutant structures, likely due to conformational heterogeneity. The electron density corresponding to the loop region (L) is only discernible to trace the main chain between TM2 and the cytoplasmic helix, but insufficient to support structural modeling. A helical model was fit into the corresponding density as a rigid body, but the registration could not be assigned due to the lack of side-chain density. Consequently, these details are omitted from the final atomic models. Interestingly, the cytoplasmic loop L and CTD are not conserved in sequence. Whereas *Mt*MscL feature a CTD juxtaposed with the transmembrane domain, neither CTD nor cytoplasmic loop are present in *Ma*MscL. Structurally, the distance between CTD and the transmembrane domain is much greater in *Ec*MscL than in *Mt*MscL. Despite the divergent sequence, the helical bundle may be structurally conserved in certain species. Our ssNMR analyses provide supplementary structural information for the CTD, suggesting the CTD helices are formed by residues 117-136 as will be presented in detail in the subsequent section.

### Solid-state NMR investigation of wild-type *Ec*MscL in liposomes

Both ^13^C and ^1^H-detected ssNMR experiments were performed on WT *Ec*MscL. ^13^C-detected experiments on uniformly ^13^C,^15^N-labeled samples enabled robust resonance assignments, while ^1^H-detected experiments employed perdeuterated ^2^H, ^13^C, ^15^N labeling to reduce dipolar broadening and improve spectral resolution. Note that due to the details of the sample production, i.e. expression in D_2_O with subsequent D/H back-exchange, protected TM regions are not expected to be NMR-visible in ^1^H-detected experiments on deuterated samples. In contrast, we expect to observe in principle all residues that are not too flexible in ^13^C-detected experiments on protonated samples, including the TM regions. Together, these complementary approaches leverage the sensitivity of ^1^H detection and the assignment reliability of ^13^C detection. For the protonated WT *Ec*MscL, we assigned 57 out of 136 residues (43%, fig. S5), predominantly located in the periplasmic loop, TM1, and CTD regions, fewer resonances are visible from TM2 and no resonances from the NTD (Fig. S6). All chemical shifts resulting from ^13^C detection can be found in Table S1. The lack of assigned resonances in TM2 and the NTD point towards structural heterogeneity in this lipid-facing region of the channel.

For deuterated WT *Ec*MscL, residue assignments were achieved for 39 out of the 136 residues (30%), with the majority of assigned signals originating from the periplasmic loop and the C-terminal domain. Within the transmembrane domains (TM1, TM2), only a few residues close to the periplasmic loop were observed with ssNMR, presumably due to limited accessibility to D/H exchange within the membrane-embedded regions. Based on these observations, we infer that the channel stays in a closed conformation during the purification and reconstitution process, as an open pore would be expected to give rise to additional signals from the pore-facing residues. The chemical shifts of all assigned nuclei of deuterated *Ec*MscL are listed in Table S2. An example of a sequential walk from T116 to E119 is shown in fig. S7.

The 2D hNH spectrum of deuterated WT *Ec*MscL (Fig. 3A) displays signals from solvent-exposed residues. Figure 3B shows the 2D NCA plane extracted from the high-resolution 3D hCANH spectrum with the assigned residues. For the deuterated WT *Ec*MscL, 85 % of the detected peaks could be assigned. The remaining 15 % are unassigned primarily due to limited sample sensitivity and inability to establish continuous backbone walks for unambiguous assignments. Nevertheless, the peak positions observed in the ^13^C-detected NCA experiment coincide well with those in the 2D NCA plane of the 3D hCANH ^1^H-detected experiment, indicating high sample reproducibility (fig. S6).

**Fig. 3.**
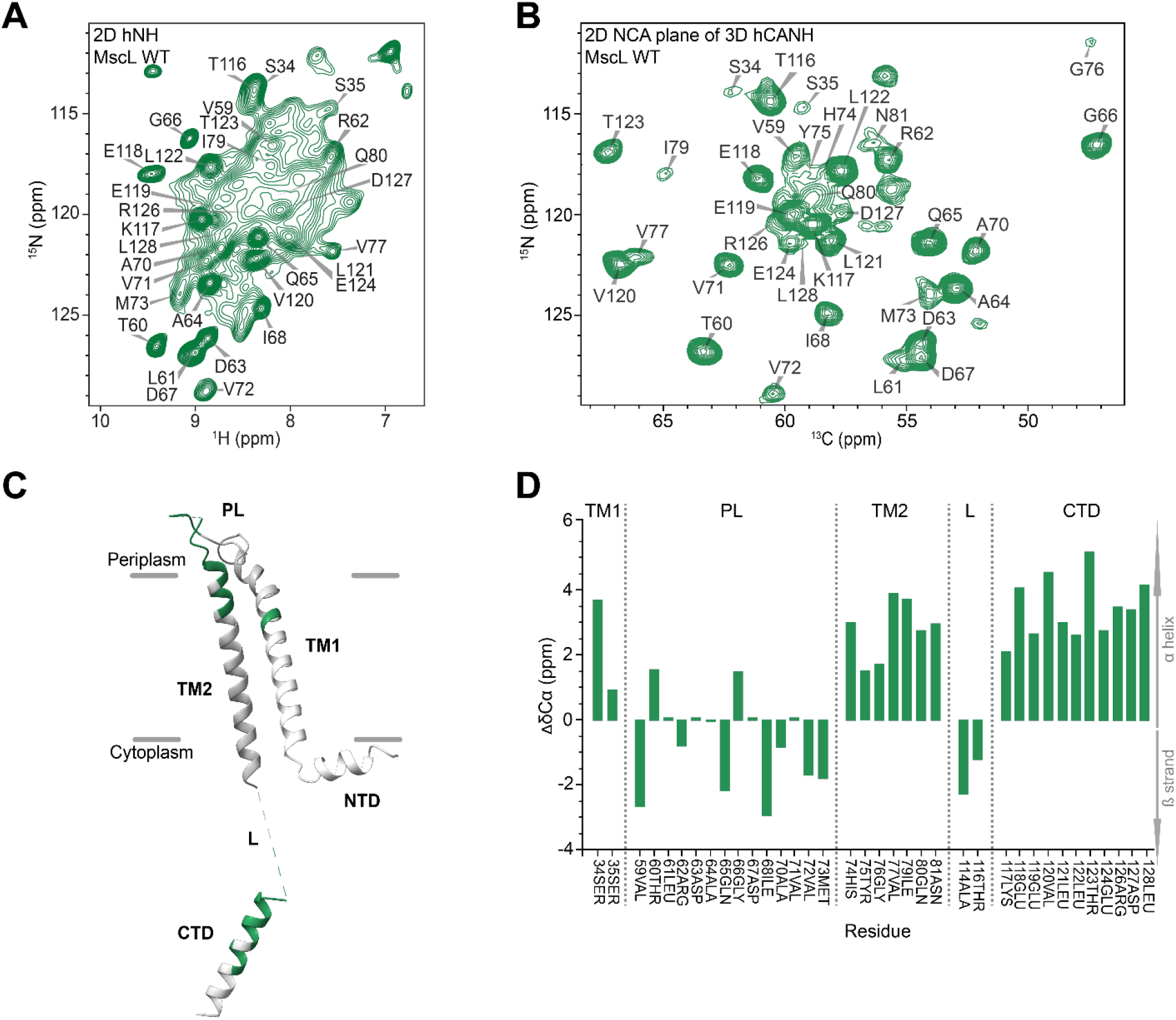
Amino acid assignments of [^2^H, ^13^C, ^15^N]-labeled wild type (WT) MscL in Azolectin liposomes by solid-state NMR. (**A**) 2D hNH ssNMR spectrum of WT MscL. (**B**) 2D NCA plane of 3D hCANH of WT MscL. (**C**) Assignments of WT MscL mapped onto a monomer of the “closed” cryo-EM structure determined here (PDB ID: 9TIU). PL = periplasmic loop, TM = transmembrane helix, L = loop, NTD = N-terminal domain, CTD = C-terminal domain. Most residues from the transmembrane regions are not visible in the ssNMR experiments, presumably due to lack of back exchange (D/H) in this area. (**D**) Secondary chemical shifts (SCS) of WT MscL (ΔδCα), shown as a function of the residues assigned. Positive SCS values indicate amino acid residues positioned within α-helical regions, whereas SCS values below zero correspond to residues located in β-strand regions.

Overall, the observed chemical shifts are in good agreement with the expected residue positions based on the cryo-EM structure of the “closed” MscL (Fig. 3C), and the secondary chemical shift analysis of WT *Ec*MscL as a function of residue number shown in Figure 3D. Positive secondary chemical shifts observed for TM1, TM2, and the CTD indicate that these regions predominantly adopt α-helical conformation.

Interestingly, residues T60 and G66, which are predicted to reside in a β-strand and the periplasmic loop, respectively, exhibit positive secondary chemical shifts in the ssNMR data, indicating a preference for α-helical conformation. This deviation may reflect the intrinsic flexibility of the periplasmic region, which could allow transient adoption of α-helical structure. Furthermore, while the “L” loop was not amenable to a cryo-EM analysis, parts of it are clearly visible in the ssNMR data. Residues A114 and T116 appear to be located in an extended conformation, as judged by their secondary chemical shifts.

### Comparison of WT and G22S *Ec*MscL structures reveals a consistently closed channel conformation in lipid nanodiscs

As discussed previously, we solved structures of both WT and the G22S mutant of *Ec*MscL in this study. Given that the G22S mutation is known to lower the threshold for channel opening in both in *in vivo* and *in vitro* functional assays (*48, 49*), we initially anticipated that the G22S mutant may increase the chance for capturing the open state conformation. However, a structural comparison of the WT MscL monomer with the G22S MscL monomer yielded a Cα RMSD of 0.83 Å over 91 pruned residue pairs, indicating a high degree of structural similarity, which is also evident from visual inspection (fig. S4D). When all 123 aligned residue pairs were included, the RMSD increased to 2.4 Å, suggesting that differences are primarily localized to flexible or loop regions. These localized structural variations are consistent with the ssNMR data (see below), where the G22S mutant exhibits increased dynamics, particularly in the periplasmic region (Fig. 5B and D).

To further assess the conformational state, the pore radius of WT *Ec*MscL was calculated using the program HOLE (*50*). The narrowest constriction is located at V23 (Fig. 2D), near the center of TM1, with a radius of around 2 Å, indicating a closed state (fig. S4). Similarly, the G22S mutant exhibits a pore profile as the wild type, consistent with a closed conformation (Fig. 2D), as the open state of MscL has been estimated to exhibit a pore diameter of approximately 30 Å (*51*–*55*). The slight differences towards the cytoplasmic side are around the hinge residue G14. Consistently, electron density corresponding to lipid tails was observed surrounding a hydrophobic pocket formed by TM2 in the G22S mutant (Fig. 4C). This pocket is defined by residues F83, L86, and F90, and corresponds to the position of L89 in *Mt*MscL (*56*). It has been proposed that channel opening requires the exclusion of membrane lipids from this pocket for the *Ec*MscS channel, where lipid interactions stabilize the closed conformation and lipid displacement is necessary for opening (*20, 57*). In WT the absence or presence of a lipid binding pocket was difficult to assess due the lower resolution obtained here as compared to the structure of G22S.

**Fig. 4.**
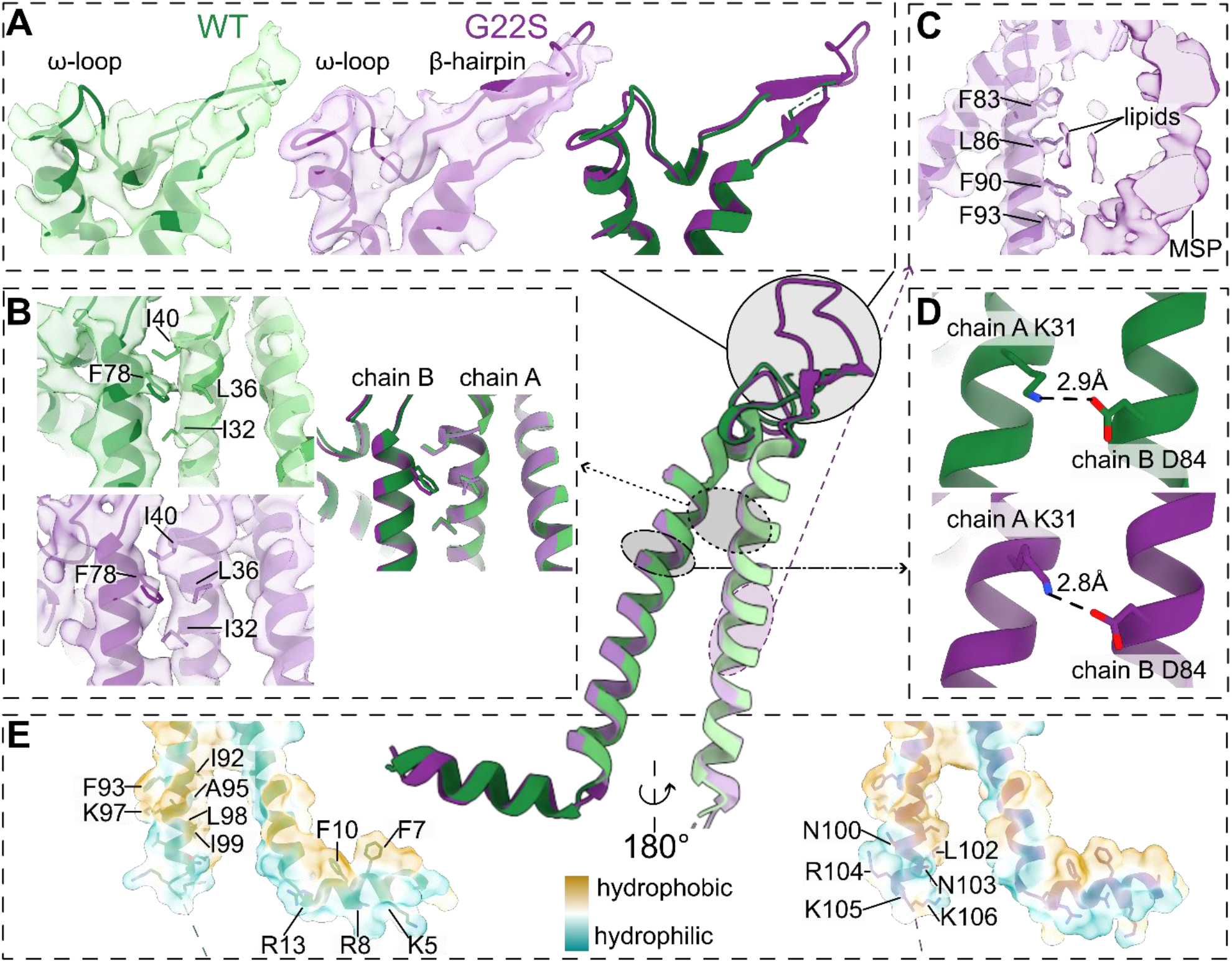
A focused view of regions of interest in the cryo-EM structures of both WT and G22S mutant of MscL. (**A**) A comparison of the periplasmic loop region (PL), which is less defined in the wild type but can be observed in ssNMR. (WT, Green; G22S, purple) (**B**) Hydrophobic pocket formed by adjacent monomer chains for F78 residues, showing no significant differences between WT and G22S structures, indicating both are in the same (closed) state. (**C**) Hydrophobic residues and proposed faint lipid density around the center of TM2. (**D**) Salt bridge formation between monomers, specifically K31 (chain A) and D84 (chain B). Both WT and G22S mutant show similar bond lengths, indicating G22S is also in the closed state. (**E**) Clustering of hydrophobic and charged residues in an amphipathic helix motif close to the membrane at the N-terminus (NTD) as well as in the cytoplasmic end of TM2.

Furthermore, it has been proposed that MscL channel opening requires substantial helical rearrangements leading to an iris-like expansion (*58*). However, in both structures obtained in this work, the pentamer remains tightly associated through a combination of van der Waals interactions at the periplasmic side and specific inter-subunit salt bridges (Fig. 4D). One example is the K31-D84 salt bridge, which exhibits distances of 2.9 Å and 2.8 Å in WT and G22S, respectively. The preservation of this salt bridge in the G22S mutant indicates minimal conformational changes in the periplasmic region. In addition to these electrostatic interactions, monomer association is further stabilized by hydrophobic contacts, notably involving F78, which inserts into a pocket formed by I32, L36, and I40 of a neighboring TM1 helix (Fig. 4B). The salt bridge as well as the hydrophobic pocket have been previously shown to be essential for maintaining the closed channel conformation (*59*).

### Solid-state NMR detects enhanced conformational dynamics in the *Ec*MscL G22S mutant

Notably, ssNMR reveals pronounced conformational differences in the G22S mutant, as evident from the overlay of the 2D NCA planes extracted from the 3D hCANH spectra of G22S and WT *Ec*MscL (Fig. 5A). For the G22S sample, 27 out of 136 residues were assigned (Table S3). While a subset of peaks can be assigned to the same residues in both spectra, many exhibit significant chemical shift perturbations, as well as the appearance or disappearance of signals between the two variants. Additionally, comparison of the 2D hNH spectra of WT (900 MHz) and G22S (600 MHz) in Fig. 5B shows substantial line broadening in the G22S mutant that cannot be explained by the different fields used alone. This broadening likely reflects increased conformational dynamics and heterogeneity, as line broadening is indicative of motions on the microsecond time-scale (*60*). Consistently, such enhanced dynamics align with the expected increase in conformational sampling in the G22S mutant, where elevated flexibility lowers the energy barrier for gating (*39*). Mapping the assigned residues of WT and G22S *Ec*MscL onto the cryo-EM structure further highlights these differences (Fig. 5C). The most striking feature is the absence of signals from the periplasmic region (residues 59-61 and 64-72) in the G22S mutant, which are clearly observed in the WT (green). This loss of detectable resonances correlates with reduced local resolution in the corresponding cryo-EM structure, supporting increased flexibility of the periplasmic region in the mutant (Fig. 4A).

**Fig. 5.**
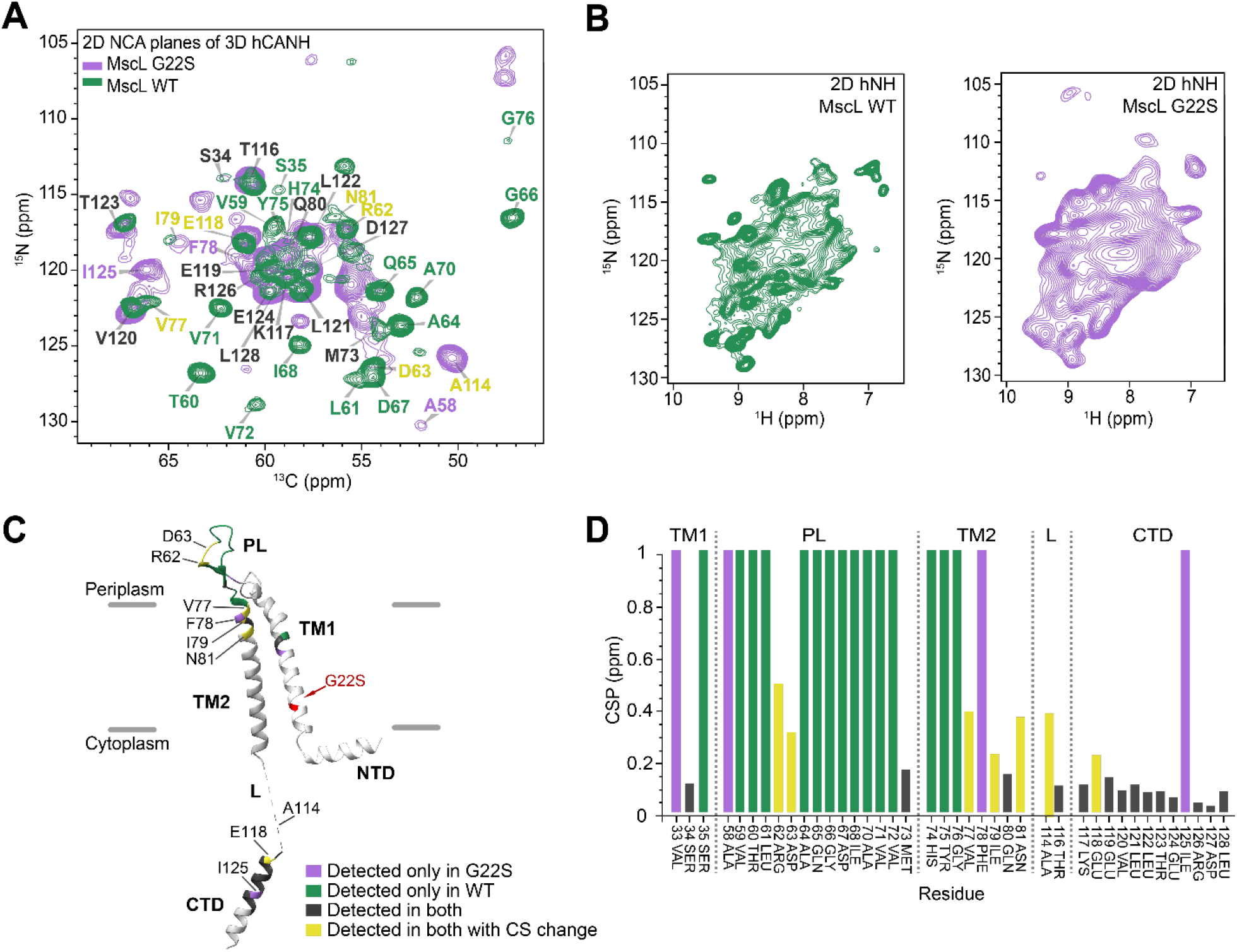
ssNMR detects G22S-induced changes in MscL. (**A**) 2D NCA plane of 3D hCANH spectrum (900 MHz, 60 kHz MAS) of WT MscL (green) overlaid with a corresponding 2D NCA plane (600 MHz, 55 kHz MAS) of the G22S mutant (purple). Black labels represent peaks observed in both spectra with similar chemical shifts. Purple labels indicate peaks only observed in the spectra of the G22S mutant. Green labels represent peaks only observed in spectra of WT MscL. Yellow labels indicate those peaks observed in both samples where the chemical shift perturbations (CSPs) are larger than 0.2 ppm. (**B**) Comparison of 2D hNH spectra of WT MscL (green, 900 MHz) and the G22S mutant (purple, 600 MHz). (**C**) Assignments of G22S MscL mapped onto a monomer of the corresponding mutant MscL cryo-EM structure, using the same criteria as in (A). (**D**) Chemical shift perturbations between WT and G22S MscL plotted as a function of residue. The color coding is the same as in (A). Note, that for residues only observed for either WT or mutant, a value of “1” is plotted.

We further analyzed the chemical shift perturbations between G22S and WT *Ec*MscL, which report on residue-wise changes in the local chemical environment (Fig. 3D). Beyond the periplasmic region, additional alterations are evident in other parts of the protein. Notably, residues R62 and D63 remain detectable in the G22S mutant but exhibit substantial chemical shift changes, while signals from the surrounding region largely disappears. At the same time, new peaks emerge in the G22S spectra (purple), particularly F78, a residue previously identified as a key tension sensor critical for channel gating (*59*). F78 resides in the TM2 region, where several other residues also exhibit substantial perturbations (see also Fig. 4B). The largest chemical shift differences (yellow; >0.2 ppm) are observed for R62, D63, V77, I79, N81, A114, and E118. The shift of E118 is consistent with previous MD simulations and patch-clamp studies, which identified this residue as undergoing the largest mobility changes during gating (*61*). In contrast, residues 120-128 in the cytosolic domain remain largely unchanged in both mutant and WT, in agreement with MD results indicating that the CTD is the most stable part and functions as a molecular sieve during channel opening (*61, 62*). Notably, I125 from this otherwise stable region is only visible in the G22S mutant, suggesting subtle differences in CTD dynamics between G22S and WT *Ec*MscL.

## Discussion

### A combination of cryo-EM and ssNMR reveals the structure of *Ec*MscL in native-like membranes

MscL is an ion channel that is gated solely by membrane tension and a foundational model system for studying how mechanical forces are transduced into protein changes at the molecular level. Several structures of MscL have been reported, most notably for *Mt*MscL and *Ma*MscL, determined in detergent micelles. Beyond these crystallographic structures, a recent ssNMR study of *Ma*MscL in both synthetic and native-like bilayers provided several distance restraints that enabled a refined structure of *Ma*MscL in a membrane environment (*38*–*40*). Despite these structures and the relatively high sequence conservation among *Mt-Ma-*, and *Ec*MscL (fig. S2), no structure of *Ec*MscL been reported to date. Early ssNMR studies of protonated *Ec*MscL using ^13^C-detected experiments mainly established the feasibility of investigating this system, but did not include detailed spectral analysis or residue-specific assignments (*37*).

Here, we present the first high-resolution cryo-EM structure of wild-type *Ec*MscL in MSP-based nanodiscs (3.7 Å, Fig. 2A). The structure is consistent with secondary structure predictions from the amino acid sequence and agrees well with previous homologous MscL structures. In parallel, we achieved residue-specific assignments for 42% (57 out of 136) of the residues of WT *Ec*MscL in native-like liposomes using ssNMR. These assignments are predominantly derived from the periplasmic loop, TM1, and the C-terminal domain, reflecting either limited hydrogen-deuterium back exchange in the case of ^1^H detected experiments or increased mobility in the case of ^13^C detected methods on protonated samples that do not rely on H/D back-exchange. The latter aspect is relevant for the lack of assignments in TM2 and the N-terminal domain. We note that this agrees well with previous solid-state NMR results on *Ma*MscL. Thus, increased dynamics of MscL NTD and TM2 appear as a universal characteristic in the context of liposomes.

In addition to the wild-type structure we have in this work also determined the cryo-EM structure of G22S *Ec*MscL, in which the mutation leads to a channel that is easier to be opened by tension (*48*). While the structure appears similar to the wild-type structure, improved sample optimization allowed us to obtain the structure of G22S at slightly higher resolution (3.5 Å, Fig. 2 and fig. S3). This in turn allowed us to characterize in more detail the PL region as well as binding of lipids into a specific binding pocket.

Previous work on *Ec*MscL (*63*) and *Mt*MscL (*64*) has emphasized the conserved nature and structural importance of the CTD, although its precise role in gating remains unresolved. The CTD is the most conserved region and is proposed to function both as a molecular sieve and as a stabilizing element of the channel (*61*). Notably, this domain remained undetected in earlier ssNMR studies of *Ma*MscL (*38*–*41*), whereas our data now enabled the assignment of 15 out of 27 CTD residues in *Ec*MscL.

### The closed state of the G22S mutant

Comparison of the *Ec*MscL structures presented here with homologous closed-state structures of *Mt*MscL (*12*) and *Ma*MscL (*46*), as well as the reported “expanded” *Ma*MscL structure (*46*), indicates that both WT and G22S mutant structures represent the closed state, as channel opening requires substantial rearrangement of the transmembrane domains and the formation of a large pore. Previous studies, including work on the related mechanosensitive channel of small conductance (MscS) and its homologs, have proposed gating mechanisms such as iris-like expansion and asymmetric or corkscrew-like motions of TM1 (*62, 65, 66*). To probe these conformational transitions, we characterized the G22S mutant alongside the WT, aiming to capture both closed and open states. G22S was chosen based on prior studies showing a reduced energy barrier for channel opening, while maintaining sufficient *E. coli* growth for sample production (*48*). However, cryo-EM densities for WT and G22S in peptide-based lipid nanodiscs show no major global differences, with only subtle local variations in the central pore and the more flexible loop regions. In contrast, the differences are more evident in the solid-state NMR investigations performed on MscL incorporated into liposomes. Signals corresponding to the periplasmic loop in WT are absent in the G22S sample, reflecting increased flexibility and highlighting this region as highly mobile in the mutant. Notably, even in the WT, chemical shifts of T60 and G66 suggest α-helical character, whereas the cryo-EM density in this region appears as β-strands and loops (Fig. 4A). This discrepancy likely reflects intrinsic flexibility, allowing this segment to sample intermediate conformations at room temperature.

Additionally, the small chemical shift differences in the CTD between WT and G22S indicate that the C-terminus remains largely stable in both variants. However, two residues in this region - E118 and I125 - show notable perturbations. Previous studies of various MscL homologs have reported that upon channel opening, only the upper part (i.e. more close to the membrane) of the CTD bundle (residues A110 to ∼E119) partially dissociates or bends outward (*61*), which may account for the observed shifts of A114 (“L” region) and E118 in G22S relative to WT. Notably, the appearance of the I125 resonance in the G22S spectra, but not in WT MscL, suggests reduced local dynamics of this residue in the mutant.

Taken together with the global similarity of the cryo-EM structures, these data indicate that the G22S mutant, despite its enhanced dynamics, predominantly adopts a closed-state in both cryo-EM and ssNMR samples. While ssNMR reveals clear differences between G22S and WT, these are not consistent with the large-scale rearrangements expected for channel opening. Instead, we propose that in the liposome environment that is less constrained than the peptide-based nanodiscs used for cryo-EM, the channels have more degrees of freedom to move towards partially open conformations, yet not fully reaching a completely open state without applying further measures. Another possible scenario is that in the G22S mutant, channels open and close with time, however stay mostly in the closed conformation.

### MscL channel opening is inhibited by tight helical packing

The gating of MscL was proposed to be driven by membrane thinning under tension, which induces tilting of the transmembrane helices and leads to an iris-like expansion while preserving helical integrity. Early mutational studies (*5, 67, 68*) and comparison with the expanded *Ma*MscL structure (PDB ID: 4Y7J), related MscS-channel studies, as well as the here determined AF3-based model of G22S *Ec*MscL (fig.S1D), support this mechanism, consistent with computational predictions (*17, 69*–*71*). Together, these conformational changes imply a substantial energy barrier, explaining why the MscL channel opens only under extreme membrane tension.

To prevent undesired spontaneous activation, several structural features stabilize the closed state. Tight helical packing, mediated by small hydrophobic residues at the pore constriction (e.g., L17 and V21 in *Mt*MscL, V23 in *Ec*MscL), forms a hydrophobic gate that resists water penetration and increases the energetic cost of helix tilting (fig. S4) (*56, 72*). In particular, the hydrophobicity of residue 22 in *Ec*MscL modulates the activation barrier: more hydrophobic substitutions raise it, whereas more hydrophilic substitutions lower it (*48*). Additional regulatory regions identified by mutational studies (*2*), can be broadly classified into inhibitory and modulatory elements, as summarized in Fig. 6. Inhibitory regions primarily stabilize tight TM1-TM2 packing in the closed state. Key elements include an inter-monomer hydrophobic interface, centered on F78 interacting with I32, L36, and I40 of a neighboring subunit and a salt bridge between K31 and D84 (Fig4. B and D). The salt bridge tethers TM1 of one subunit to TM2 of the adjacent subunit, contributing to the energetic barrier for opening (*59*). Disruption of the F78-centered hydrophobic pocket further increases this energetic cost and may also help maintain proper helical packing of the pentamer in the closed state. Consistently, the loss-of-function mutant F78N underscores the importance of this interaction in stabilizing the closed-state conformation (*59, 62*). In our structures, F78 occupies a similar hydrophobic pocket in both WT and G22S. However, F78 is only observed in the G22S ssNMR spectra, suggesting that it is partially exposed to water but rigid enough to be observed in NMR. In contrast in WT, inaccessibility to H/D exchange and thus to back protonation makes it NMR invisible. Possibly, in the G22S mutant, the protein temporarily opens – while being mostly in the closed state – and during these opening events, F78 leaves the hydrophobic pocket and can thus become protonated. In summary, although we did not capture an open-state conformation of *Ec*MscL, the ssNMR data provide the first experimental glimpse into conformational changes that shift the channel from the closed-state towards the open state.

**Fig. 6.**
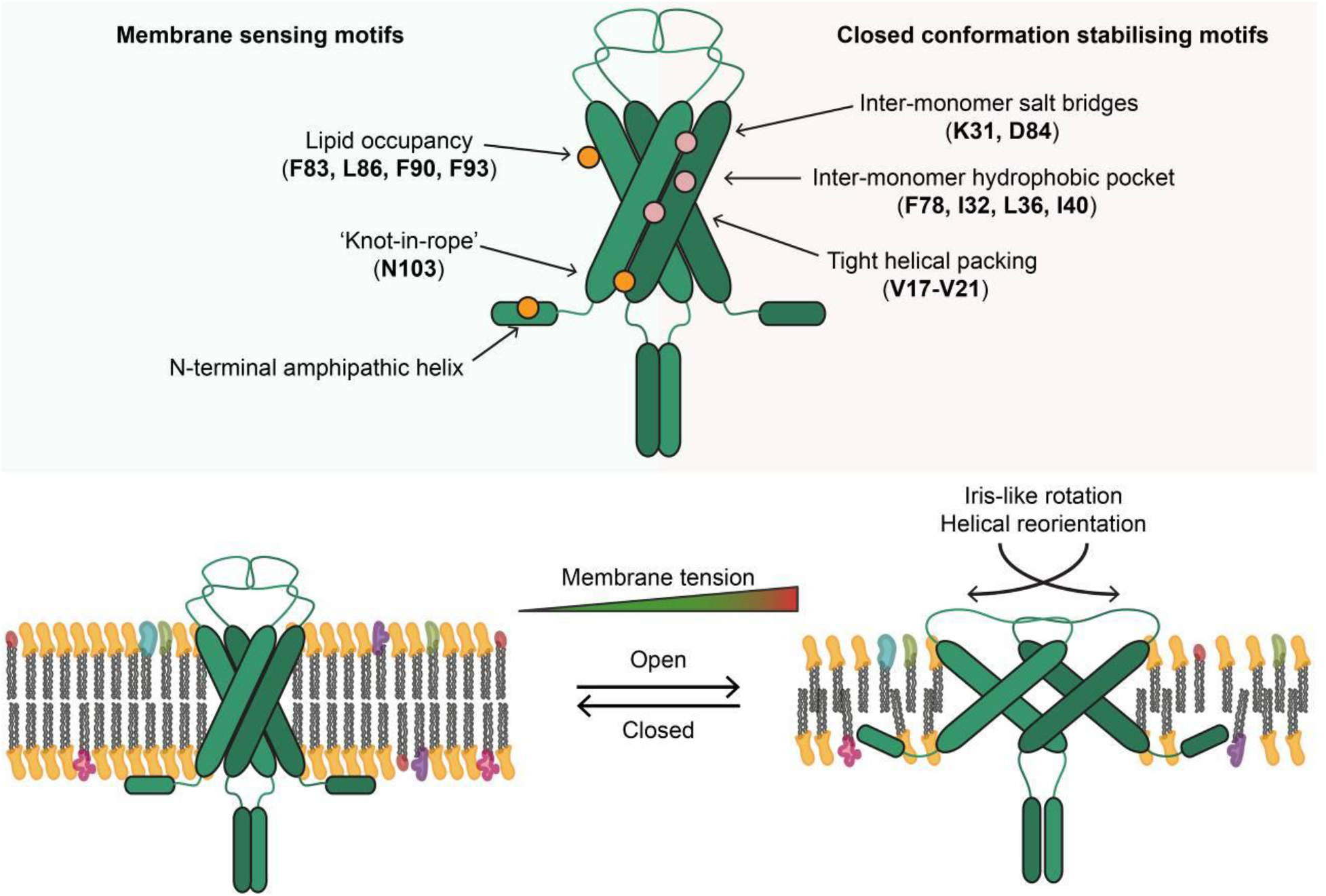
Proposed mechanism of channel regulation by *Ec*MscL. Membrane sensing motifs (in orange) sense an increase in membrane tension by a reduction in lipid occupancy, increased unit area per lipid, and decreased hydrophobic thickness. The closed-state conformation of MscL is stabilized by several helical-associating motifs (shown in magenta), most notably inter-monomer salt bridges and an inter-monomer hydrophobic pocket, as well as general tight-helical bundling.

### MscL opening is triggered by the membrane properties

Regulation of MscL opening is closely linked to membrane tension and associated changes in the membrane. It has been shown that purified mechanosensitive channels such as *Ec*MscS and *At*MSL10 (*73*) adopt open conformations upon delipidation, whereas reintroduction of lipids restores the closed state of *Mt*MscL (*56*) and *Ec*MscS (*74*–*76*). In MscS, lipids occupying hydrophobic pockets between transmembrane helices stabilize the closed state via interactions with the acyl core of the membrane(*77, 78*), a mechanism likely conserved in MscL, which possesses a similar TM pocket. In our structures, putative lipid densities were observed near the TM2 region of *Ec*MscL (Fig. 4C). Although no specific interactions could be observed, the density is indicative of a semi-stable lipid association in this region. Supporting this, previous MD simulations of *Mt*MscL show that the lipid acyl chains occupy a cavity formed by I82, V86, and V22 (*62*). These simulations further suggest that channel opening is partially coupled to lipid rearrangement, where reduced lipid packing exert outward forces on TM2, promoting channel expansion.

The N-terminal helix exhibits a sequence and spacing characteristic of an amphipathic helix (1-14 MSIIKEFREFAMRG) (Fig. 4E). The positively charged residues can interact with negatively charged lipid headgroups, while hydrophobic side-chains further interact with the lipid tails (*47, 79*). Amphipathic helices are known to sense changes in lipid packing and membrane curvature (*80*–*82*), consistent with a potential role of the N-terminus in mechano-sensing. In line with this, our cryo-EM structures of both WT and G22S *Ec*MscL show membrane association of the N-terminal region (Fig. 2). However, how signals from this amphipathic helix are transmitted to the transmembrane core remains unclear and warrants further investigation.

Mutational studies have further indicated that N103 acts as a “knot in the rope”, preventing charged residues at the cytoplasmic end of TM2 from penetrating the membrane core (*83, 84*). In our structure, at higher map contour levels, N103 of the WT channel exhibits extra density connecting to the adjacent N-terminus, suggesting a stabilizing role in helix-helix interaction in the closed state and restricting helix displacement within the membrane. Such a charged “knot” that anchors and modulates transmembrane tilt and gating motions may represent a general regulatory principle across channels, including K2P channels such as TREK-1 and TREK-2 (*85*).

## Conclusion

Here, we have solved the cryo-EM structures of wild-type and G22S MscL from *E. coli*, which is a model system for mechanosensitive channels. In the lipid nanodiscs used for cryo-EM, both wild-type and G22S adopt a closed-state conformation. In liposomes as used for solid-state NMR, the G22S mutant which opens at lower membrane tension, already experiences a shift towards the open state, however still remains in the closed state predominantly. This is seen by significant chemical shift differences and the appearance and disappearance of signals upon the G22 mutation, e.g. the F78 gate keeper residue that is only visible in the G22 mutant. Large changes are also observed for the periplasmic loop that plays a major role in channel opening. From a methodological point of view it is important to note that only the combination of cryo-EM and solid-state NMR allowed for a comprehensive understanding of the structural and dynamic effects of the G22 mutant. The open state structure of G22 predicted by AlphaFold3 provides another indication of how the open state of MscL will look like. Future work will aim at determining the open state structures of wild-type or G22S MscL experimentally. From a technical point we note that the structure of MscL in lipid nanodiscs was obtained without further stabilizing or enlarging elements such as nanobodies or protein modification such as designed helical repeats and it thus represent a technically demanding structure determination with sufficiently high resolution. At the same time the solid-state NMR data quality of MscL (both WT and G22S) was excellent, in particular if compared to other membrane proteins reconstituted into liposomes. The data presented here will thus be an excellent basis for future work that focuses on the open state, the transition between the closed and the open states, and the dynamics of the protein with quantitative relaxation based solid-state NMR measurements.

## Materials and Methods

### Expression and purification of MSP1E3D1

MSP1E3D1 with a tobacco etch virus (TEV) protease-cleavable N-terminal 7×His tag was expressed in *E. coli* BL21(DE3) transformed with the plasmid pMSP1E3D1, which was a generous gift from Stephen Sligar (Addgene plasmid #20066; http://n2t.net/addgene:20066; RRID: Addgene_20066). Freshly transformed BL21(DE3) cells were grown in terrific broth (TB) medium with 50 μg/ml kanamycin at 37 °C. Protein expression was induced at an OD_600_ of around 0.6-0.8 by addition of isopropyl β-d-1-thiogalactopyranoside (IPTG) to a final concentration of 1 mM. Bacteria were harvested after 4 h expression at 37 °C by centrifugation at 5,000 *g* for 30 min at 4 °C. Bacterial pellets were resuspended and lysed by high-pressure homogenization at 15,000 psi using an LM10 microfluidizer (Microfluidics, USA) in PBS with 1% Triton X-100 (w/v), 1 mM PMSF. DNase and protease inhibitor (pepstatin A) were added at recommended dosage. Unsolubilised debris was removed by centrifugation at 20,000 g for 30 min at 4 °C. The resulting supernatant was loaded onto a 5 ml HisTrap HP column (GE healthcare) using an Äkta Pure25 chromatography system. The column was washed with each 5 column volumes of wash buffer I (40 mM Tris, pH 8.0, 300 mM NaCl, 1% Triton X-100 (w/v)), wash buffer II (40 mM Tris, pH 8.0, 300 mM NaCl, 50 mM sodium cholate, 20 mM imidazole), and wash buffer III (40 mM Tris, pH 8.0, 300 mM NaCl, 50 mM imidazole) in succession. His-MSP1E3D1 was eluted by 40 mM Tris, pH 8.0, 300 mM NaCl, 400 mM imidazole in 5 CVs. Protein containing fractions (as determined by Abs_280_) were pooled and digested with His-tagged TEV protease at 1:20 (w/w) during dialysis against 20 mM Tris, pH 7.4, 100 mM NaCl, 0.5 mM EDTA, at 4 °C overnight. Cleaved His-Tag and TEV protease were removed by another round of IMAC purification. Purified MSP was concentrated by Amicon Ultra 15-ml 10-kDa cut-off centrifugal filters (Millipore Sigma) to a concentration of 7 mg/mL, flash frozen in liquid nitrogen, and stored at ‐80 °C until further use.

### Expression and purification of wild-type and G22S mutant MscL

DNA fragments coding for the wild-type and G22S mutant *Escherichia coli* MscL were cloned into a pET21a vector and used to transform *E. coli* BL21(DE3) cells. Bacteria were cultivated in LB medium with 50 μg/ml carbenicillin at 37 °C until an OD_600_ of 0.8. The temperature was reduced to 25 °C and protein expression was induced by the addition of 0.5 mM IPTG. After 16 h, cells were collected by centrifugation at 4,000 g for 20 min at 4 °C. To achieve uniform ^13^C,^15^N labeling of MscL (wild type and mutant form), minimal M9 media with ^15^NH_4_Cl and ^13^C_6_-glucose (Cambridge Isotope Laboratories, USA) as the sole nitrogen and carbon sources were used instead of LB media. For the expression of perdeuterated ^13^C,^15^N labeled protein a complete media exchange had to be implemented: Freshly transformed and carbenicillin selected *E. coli* were cultivated in LB medium with 50 μg/ml carbenicillin at 37 °C and 165 rpm until the OD_600_ reached 0.8. Bacteria were gently spun down (2,000 g, 4 °C) for 10 minutes and carefully resuspended in half the initial culture volume of pre-warmed perdeuterated M9 media with ^15^ND_4_Cl and ^13^C_6_-D7-glucose (Cambridge Isotope Laboratories, USA) as the sole nitrogen and carbon sources. After incubation at 37 °C for 30 min, the temperature was lowered to 25 °C. After an additional hour, expression was induced by addition of 0.5 mM IPTG. After 16 h incubation at 25 °C and 165 rpm, bacteria were spun down at 4,000 g for 20 min at 4 °C.

Cell pellets were resuspended in PBS supplemented with 1 mM PMSF, 1 μM leupeptin, and 1 μM pepstatin, lysed by high pressure homogenization at 15,000 psi at 4 °C using an LM10 microfluidizer (Microfluidics, USA) and 3 % Triton X-100 (w/v) was added. The supernatant was clarified by centrifugation at 20,000 g for 30 min at 4 °C and incubated for 1 h at 4 °C with TALON beads (Cytiva) at a concentration of 0.5 mL of beads per liter of culture. Loaded beads were transferred to a gravity flow column and washed consecutively with 5 column volumes of MscL-washing Buffer I (PBS, 0.2% Triton X-100 (w/v), 35 mM imidazole) and MscL-washing Buffer II (20 mM HEPES, pH 7.4, 100 mM KCl, 0.2% Triton X-100 (w/v)). The protein was eluted by 20 mM HEPES, pH 7.4, 100 mM KCl, 0.2% Triton X-100 (w/v), and 300 mM imidazole and dialyzed overnight against 20 mM HEPES, pH 7.4, 100 mM KCl, 0.2% Triton X-100 (w/v) using 6-8 kDa MWCO dialysis tubing (Spectrum).

Protein concentration was estimated by a BCA assay following supplier instructions (Thermo Fisher).

### Reconstitution of labeled MscL into proteoliposomes for solid-state NMR

Soy phospholipids (azolectin, Avanti Polar Lipids, USA) were resuspended to a concentration of 20 mg/mL in 20 mM HEPES, pH 7.4, 100 mM KCl, 6% Triton X-100 (w/v) for 1 h at room temperature with rigorous agitation. Impurities and unsolubilised aggregates were removed by centrifugation for 5 minutes at 10,000 g and 4 °C. The resulting lipid suspension was added to purified (isotope-labeled) MscL at a 1:1 (w:w) ratio and dialysed against 20 mM HEPES, pH 7.4, 100 mM KCl at 4 °C. Proteoliposome formation was initiated by detergent removal: 1 g Bio-Beads (BioRad) per 1 mg Triton X-100 were added followed by 3 h incubation at room temperature under gentle agitation. Bio-Beads were removed using a disposable gravity column. Proteoliposomes were pelleted by ultracentrifugation (100,000 g, 1 h at 4 °C).

### Reconstitution of MscL into MSP-nanodiscs

All lipids were purchased from Avanti Polar Lipids (USA) and solubilized with 100 mM sodium cholate in 20 mM Tris-HCl, pH 7.4, and 150 mM NaCl with sonication. Purified MscL and MSP1E3D1 were mixed with solubilized *E. coli* total lipid extract, at an estimated molar ratio of 1:4:240 of MscL:MSP1D1E3:lipid and incubated for 1 h at room temperature with gentle agitation. In order to remove the detergent, the mixtures were incubated with 1:6 g/ml (w/v) Bio-Beads for 1.5 h without agitation and an additional 16 h under gentle agitation. After the addition of another 1:3 g/ml (w/v) Bio-Beads, samples were incubated under gentle agitation for an additional 2 h. Bio-Beads were removed by filtration, and the samples were loaded on cobalt-affinity columns (TALON) to remove empty nanodiscs. The MscL-loaded nanodiscs were eluted in 20 mM Tris-HCl, pH 7.4, 150 mM NaCl, 400 mM imidazole for wild type and in 20 mM HEPES, pH 7.4, 100 mM KCl, 400 mM imidazole for the G22S mutant. The eluted samples were then concentrated using Amicon Ultra 15-ml 50-kDa MWCO centrifugal filters (EMD Millipore/Merck) to a volume of 1 mL, and injected onto a Superdex 200 10/300 increase column (Cytiva, USA) with a flow rate of 0.5 mL/min in 20 mM Tris-HCl, pH 7.4, 150 mM NaCl for wild type and in 20 mM HEPES, pH 7.4, 100 mM KCl for the G22S mutant. The purity of fractions was analyzed by SDS-PAGE, and the concentration was determined by a BCA assay.

### Preparation of EM grids and data collection

Negative-stain electron microscopy was performed to assess sample homogeneity prior to cryo-EM acquisition using 2% (w/v) uranyl acetate.

For the cryo-EM analysis of the EcMscL WT sample, 4 µl sample at a concentration of 0.4 mg/ml were applied to glow-discharged 300 mesh R1.2/1.3 gold grids (Quantifoil Micro Tools GmbH). Grids were prepared using a Vitrobot Mark IV (Thermo Fisher Scientific) operated at 4 °C and 95% relative humidity. After an incubation time of 10 s, grids were blotted for 2 s or 4 s with a blot force of 0 and plunge-frozen in liquid ethane cooled by liquid nitrogen. The MscL G22S mutant was concentrated to 1 mg/ml (determined by NanoDrop) and applied to glow-discharged 300 mesh R1.2/1.3 gold grids (Quantifoil Micro Tools GmbH). Grids were blotted for 2.5 s with a blot force of ‐5 at 4 °C and 95% relative humidity prior to plunge-freezing.

Cryo-EM data for wild-type MscL were acquired on a Titan Krios G3i (Thermo Fisher Scientific) transmission electron microscope operated at 300 kV equipped with a K3 Summit direct electron detector and a BioQuantum energy filter (Gatan). Movies were recorded in CDS counting mode at a nominal magnification of 105,000×, corresponding to a calibrated pixel size of 0.83 Å at the specimen level. Zero-loss images were collected using an energy filter slit width of 20 eV centered on the zero-loss peak. For datasets acquired on copper and gold grids, movies were fractionated into 32 and 34 frames per micrograph, respectively. Total exposure times were 0.86 s (copper grids) and 0.91 s (gold grids), with dose rates of 69.81 e⁻/Å^2^/s and 66.1 e⁻/Å^2^/s, resulting in total accumulated doses of 60.04 e⁻/Å^2^ and 60.15 e⁻/Å^2^, respectively. A total of 18,488 movies (9,454 from copper grids and 8,994 from gold grids) were collected automatically using EPU (version 3.3, Thermo Fisher Scientific) with a defocus range of ‐1 µm to ‐2.8 µm. A second round of data collection was collected on the same electron microscope with 0.83 Å pixel size, 60.75 e⁻/Å^2^/s, and a total dose of 60.75 e^‐^/Å^2^.

Data for the MscL G22S mutant were collected using the same microscope setup. A total of 20,802 movies were recorded at a dose rate of 38.8 e⁻/Å^2^/s, with each movie acquired over 1.56 s and fractionated into 29 frames. The defocus range was set to ‐0.8 µm to ‐2.4 µm.

### Image processing

Single particle cryo-EM datasets of WT MscL were processed in Relion 5.1 (*86*) and cryoSPARC v4.6.2 (*87*). Movies were gain-normalized, motion-corrected by patch motion correction, and patch contrast transfer function (CTF) correction in cryoSPARC. Particles were automatically picked by blob picking and extracted with a box size of 360 pixels, binned to a box size of 216 pixels. Two rounds of 2D classification were used to remove junk particles, and particles from 2D classes that showed clear secondary structural features were used to generate the ab-initio model, which was followed by template picking. 592,083 unbinned particles were re-extracted, and after removing duplicates, 352,657 particles were combined.Several rounds of ab-initio models were generated with C1 symmetry, and the well-defined classes were then further refined by local refinement with the mask of the MscL protein using C5 symmetry. The density maps were sharpened by global and local CTF refinement. To further improve the map quality, two half-maps for each reconstruction following local refinement were imported into deepEMhancer (*88*) in tightTarget mode to generate the maps guiding model building.

The datasets of G22S MscL were processed in cryoSPARC v4.6.2. Movies were gain-normalized, motion-corrected by patch motion correction and defocus values were determined by patch contrast transfer function (CTF) correction. Particles were picked using blob-based autopicking and extracted with a box size of 360 pixels, binned to a box size of 216 pixels, then subjected to 3 rounds of 2D classification. 1,567,850 particles corresponding to the intact channel with different orientations were re-extracted without binning to generate the 3D initial model. 1,049,338 particles were selected and subjected to local refinement with C1 or C5 symmetry. Global and local CTF refinement were performed to further improve the resolution. Local resolution estimates were calculated in cryoSPARC. The data processing outlines are depicted in fig. S8 and fig. S9 for WT and G22S, respectively. Statistics for data collection and refinement are provided in table S4.

### EM model building and structural refinement

An initial model of *E*.*coli* MscL was obtained with Alphafold3, and the structure was then manually built into the cryo-EM density map using *Coot* 0.9.6.2 (*89*), followed by iterative real space refinement in PHENIX 1.21.2 (*90*), and manual adjustments in *Coot* and ISOLDE (*91*). For building the atomic models of the G22S channel, the structure of the wild-type channel was used as the initial model and adjusted manually in *Coot*.

The pore radius analysis was performed using HOLE 2.2.005 (*50*), and the figures were generated with ChimeraX 1.9 (*92*) and VMD (*93*).

### ssNMR spectroscopy and analysis

The ^13^C-detected spectra were recorded on a Bruker 700 MHz wide-bore spectrometer equipped with a 3.2 mm triple-resonance Efree MAS probehead. The PDSD spectrum was recorded at 11 kHz MAS with 20 ms ^13^C‐^13^C spin diffusion time. The NCA, NCO, NCACX, NCOCA, NCACO, NCOCX, NCACB, NCOCACB, and CONCA spectra were recorded at 17 kHz MAS and at sample temperature close to 0 °C following a previously published protocol (*94*). Heteronuclear magnetization transfers were achieved through cross polarization (CP) (*95*). Homonuclear (C‐C) magnetization transfers were achieved through BSH-CP (*96*) (for backbone CA-CO and CO-CA) and DREAM (*97*) (for CA-CB).

For the ^1^H-detected experiments, 100% H_2_O back-exchanged ^2^H, ^13^C, ^15^N labelled MscL WT and G22S mutant samples were packed into 1.3 mm rotors. 0.2 µL of saturated DSS (4,4-dimethyl-4-silapentane-1-sulfonic sodium salt) solution in D_2_O for locking was added prior to rotor closure. Solid-state NMR spectra of WT and G22S MscL were recorded on 900 MHz and 600 MHz Bruker spectrometers. All experiments were conducted at a sample temperature of 14 °C, calibrated according to the position of the water resonance relative to DSS.

G22S MscL spectra were recorded using a 600 MHz spectrometer (Bruker), equipped with a 1.3 mm triple channel (HCN) probehead (Bruker), at 55 kHz MAS frequency. Double sensitivity-enhanced 3D hCANH, hCONH, hCAcoNH, and hCOcaNH spectra were recorded for backbone assignment (*98, 99*). For the side-chain assignments, a triple sensitivity-enhanced 4D hCXCANH spectrum (*99, 100*) was recorded on a 900 MHz spectrometer (Bruker) equipped with a 1.3 mm 4 channel (HCND) VTX probehead.

WT MscL spectra were recorded on the 900 MHz spectrometer equipped with a 1.3 mm 4 channel VTX probehead at 60 kHz MAS frequency. Single sensitivity-enhanced 3D hCANH, hCONH (*101*) and triple sensitivity-enhanced 4D hCACONH, hCOCANH spectra were recorded for backbone assignments (*98, 99*). The 4D spectra were recorded using 5% non-uniform sampling (NUS). NUS lists were generated using http://gwagner.med.harvard.edu/intranet/hmsIST/ (*102, 103*).

Experimental details for all NMR experiments can be found in Tables S5-S8.

The 3D spectra were processed using TopSpin v4 (Bruker), all 4D spectra were reconstructed and processed using compressed sensing with the iterative soft thresholding (IST) algorithm implemented in nmrPipe (*104, 105*). CCPNMR v3.2 (*106*) and NMRFAM-Sparky (*107*) were used for the data analysis and chemical shift assignments.

^1^H-^15^N-^13^Cα chemical shift perturbations were calculated according to the following formula (*108*):

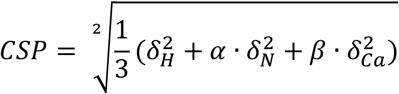

where 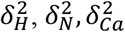 are the chemical shift differences between WT and G22S MscL, α is 0.2 for glycine and 0.14 for other amino acids, β is 0.3.

## Author contributions

T.X. and A.K. contributed equally to this study. T.X., D.Q., T.S., D.R., and S.C. obtained and analyzed single-particle cryo-EM data. A.K., C.S., C.Ö., and S.K. recorded and analyzed solid-state NMR data. T.X., C.S., and S.L. produced cryo-EM and solid-state NMR samples, respectively. A.L. designed the study. T.X., A.K., He.Sa., Ha. Su., and A.L. wrote the paper. All authors reviewed and revised the paper.

## Data and materials availability

All data are available in the main text or the supplementary materials.

