## Supplementary Materials for "Atomic structure and dynamics of the mechanosensitive channel MscL from *E. coli* by cryo-EM and solid-state NMR"

**This PDF file includes:**

Figs. S1 to S9
Tables S1 to S8

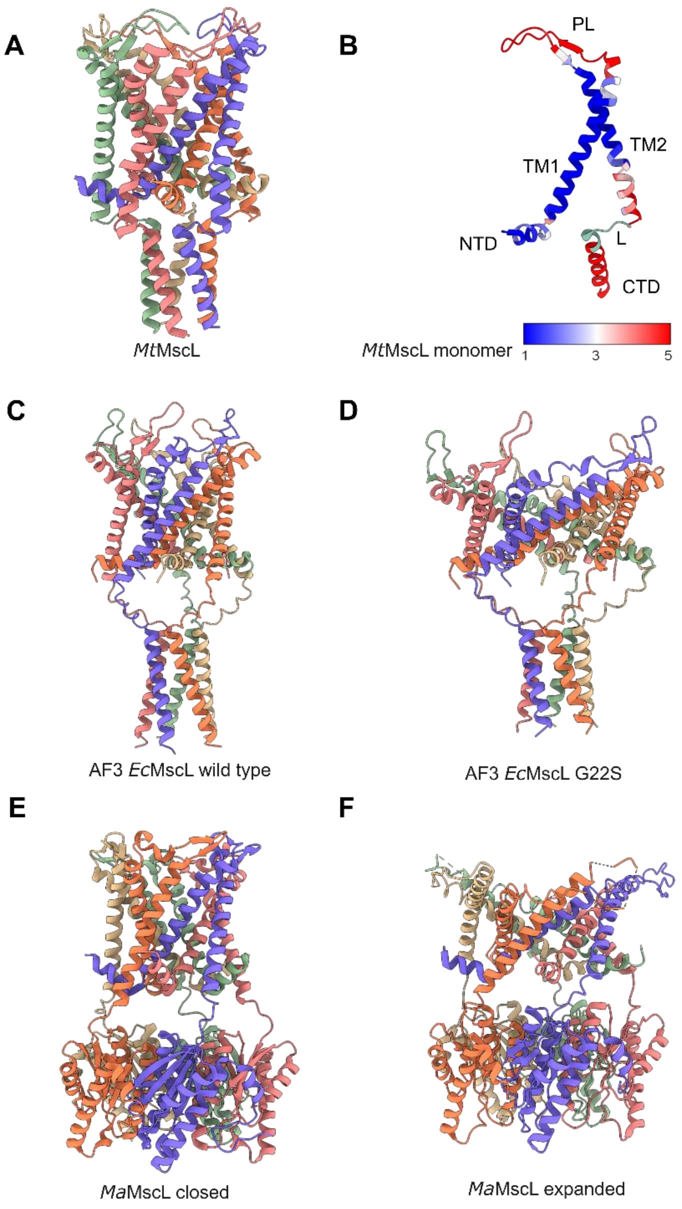


**Fig. S1. Homologous structures and Alphafold3 predictions.** (**A**) The overall structure of closed-state *Mt*MscL (PDB ID: 2OAR) from *Mycobacterium tuberculosis*. (**B**) Structural comparison of the crystal structure of *Mt*MscL (PDB ID: 2OAR, chain A) with the G22S mutant of *Ec*MscL (current study, PDB ID: 9TIV, chain A). Similarity is indicated by a red (different) to blue (similar) color scale. Missing residues (primarily in the 'L' region) are shown in green. (**C**) Alphafold3 prediction of wild-type *Ec*MscL. (**D**) Alphafold3 prediction of the *Ec*MscL G22S mutant. (**E**) The overall structure of closed-state *Ma*MscL (PDB ID: 4Y7K) from *Methanosarcina acetivorans*. (**F**) The overall structure of expanded-state *Ma*MscL (PDB ID: 4Y7J).

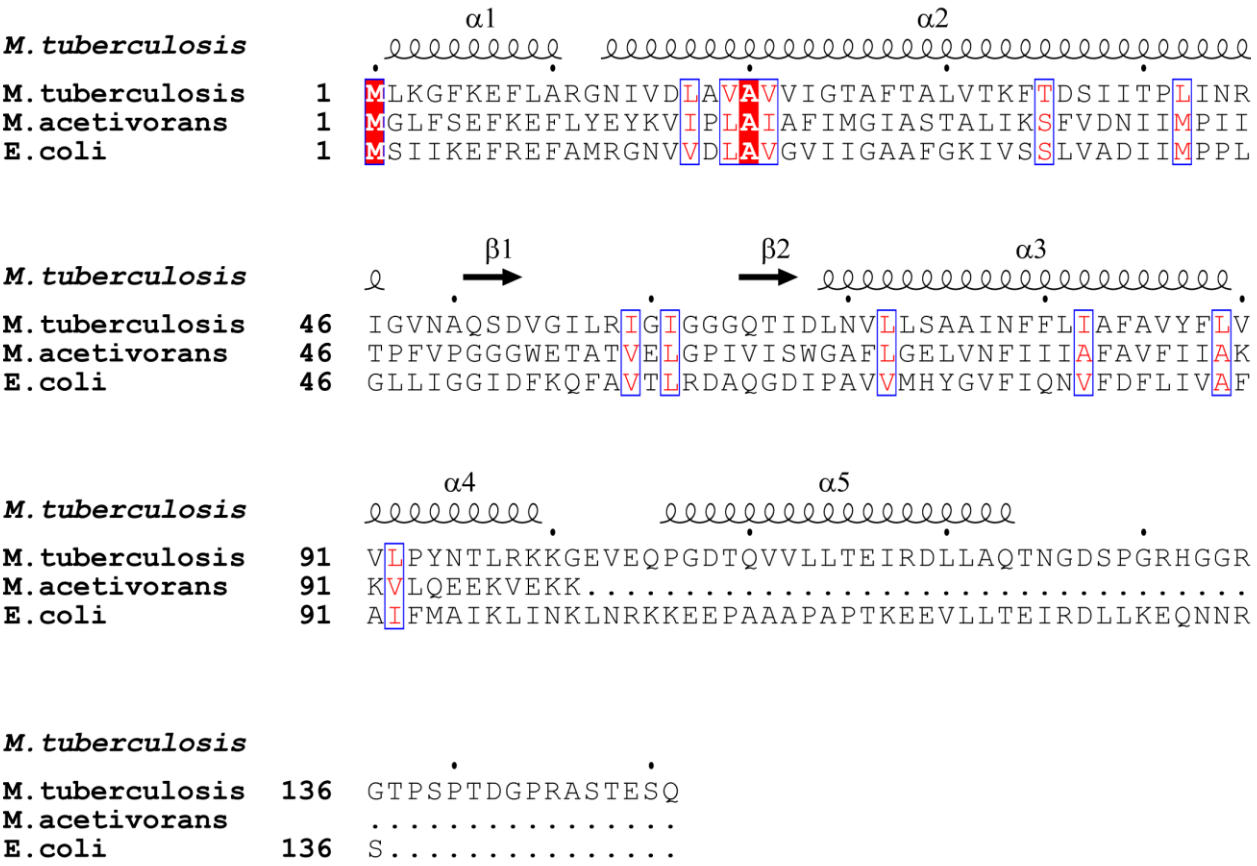


**Fig. S2. Sequence alignment of *Mt*MscL, *Ma*MscL, and *Ec*MscL by ESPript 3.2 (*109*)**

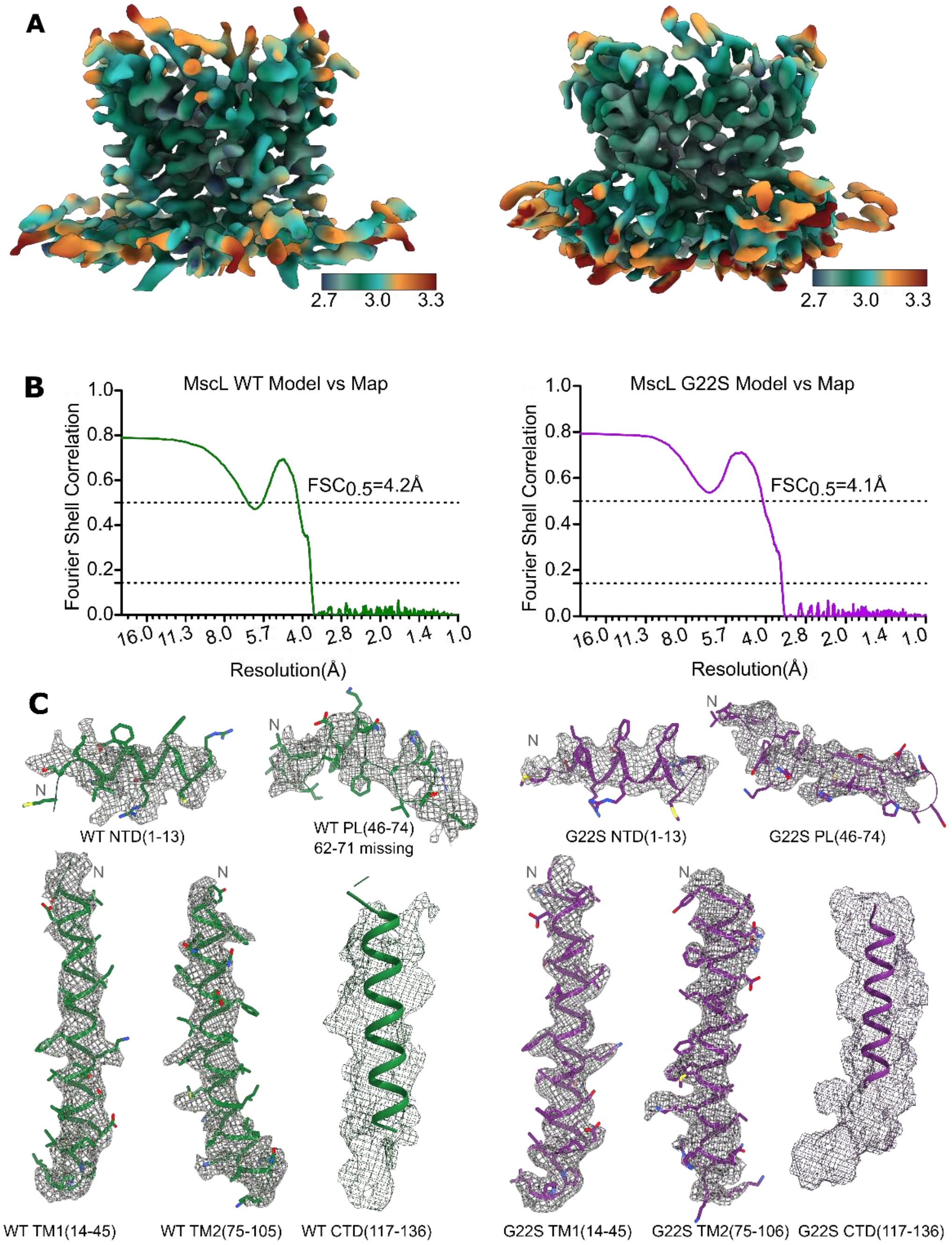


**Fig. S3. Single-particle cryo-EM characterization of MscL wild-type and mutant.** (**A**) Cryo-EM maps colored by local resolution, indicated values are in Å. (left: WT, right: G22S mutant). (**B**) FSC curves from cross validation between the atomic models against unmasked sum of both half-maps (left: WT, right: G22S mutant). (**C**) Segmented cryo-EM density maps (mesh) of WT and G22S mutant. The corresponding fitted atomic model is shown in green for WT and purple for the G22S mutant.

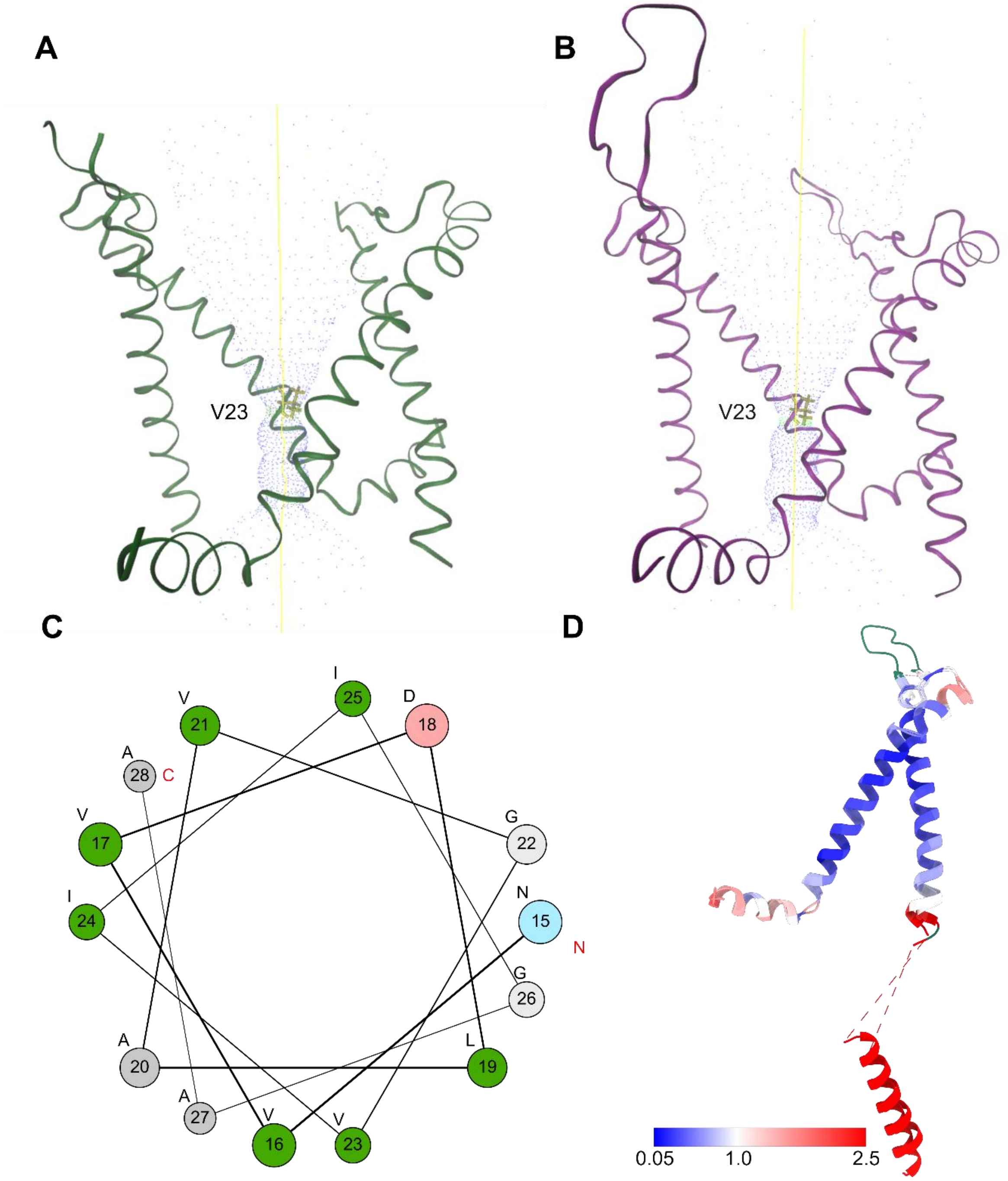


**Fig. S4. Pore radius of the *Ec*MscL channel as modelled by HOLE 2.2.005. (A)** The ion conduction pore of wild-type *Ec*MscL. Only two subunits are shown for clarity, and dots outline the central pore. The narrow position at V23 is highlighted. **(B)** The ion conduction pore of the *Ec*MscL G22S mutant. **(C)** The helical wheel from residues N15-A28. **(D)** Structural comparison of the wild-type *Ec*MscL (PDB ID: 9TIU, chain A) with the G22S mutant of *Ec*MscL (PDB ID: 9TIV, chain A). Similarity is indicated by a red (different) to blue (similar) color scale. Missing residues (primarily in the 'PL' region) are shown in green.

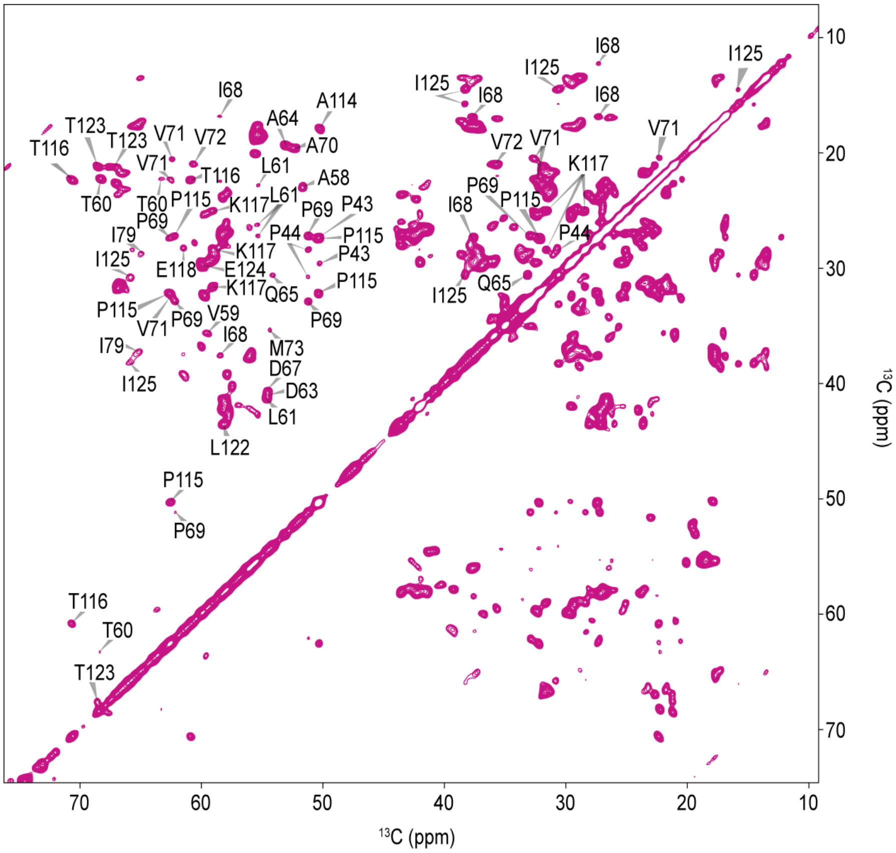


**Fig. S5. 2D $^{13}C$-$^{13}C$ PDSD spectrum.** Spectrum of [$^{13}C$, $^{15}N$]-labeled *Ec*MscL WT in Azolectin proteoliposomes and with assigned residues labelled. The spectrum was recorded on a 700 MHz spectrometer at an MAS rate of 11 kHz.

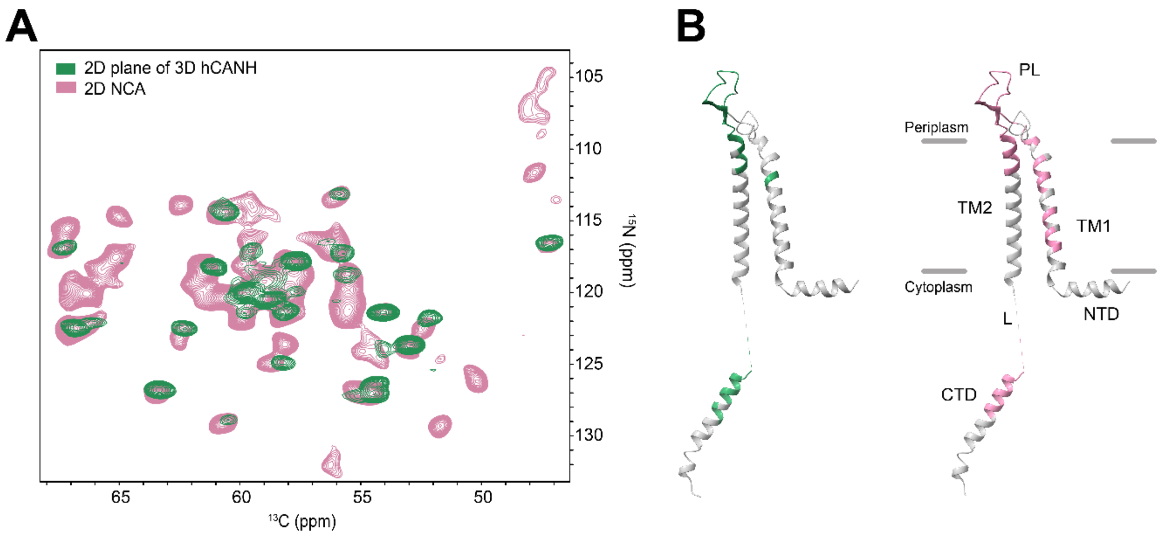


**Fig. S6. Comparison of** $^{1}$**H- and** $^{13}$**C-detected spectra.** (**A**)**.** Overlay of $^{13}$C-detected 2D NCA (3.2 mm, 700 MHz) and the 2D plane of a $^{1}$H-detected 3D hCANH spectrum (1.3 mm, 900 MHz) of *Ec*MscL WT in Azolectin. As a result of deuteration and subsequent H/D exchange, less peaks can be observed in the 2D plane of the $^{1}$H detected spectrum (green), but the presented peaks coincide with the $^{13}$C-detected 2D NCA spectrum (pink). **(B).** Assignments of WT *Ec*MscL achieved using either $^{1}$H-detected (green) or $^{13}$C-detected (pink) spectra, and mapped onto a monomer of the here determined cryo-EM structure (PDB ID: 9TIU). PL = periplasmic loop, TM = transmembrane helix, L = loop, NTD = N-terminal domain, CTD = C-terminal domain. Missing residues in TM1 and TM2 in the $^{1}$H-detected data could result from a lack of H/D back-exchange. In the case of the $^{13}$C-detected data, however, all residues are fully protonated and thus the lack of peaks for TM2 likely results from dynamics.

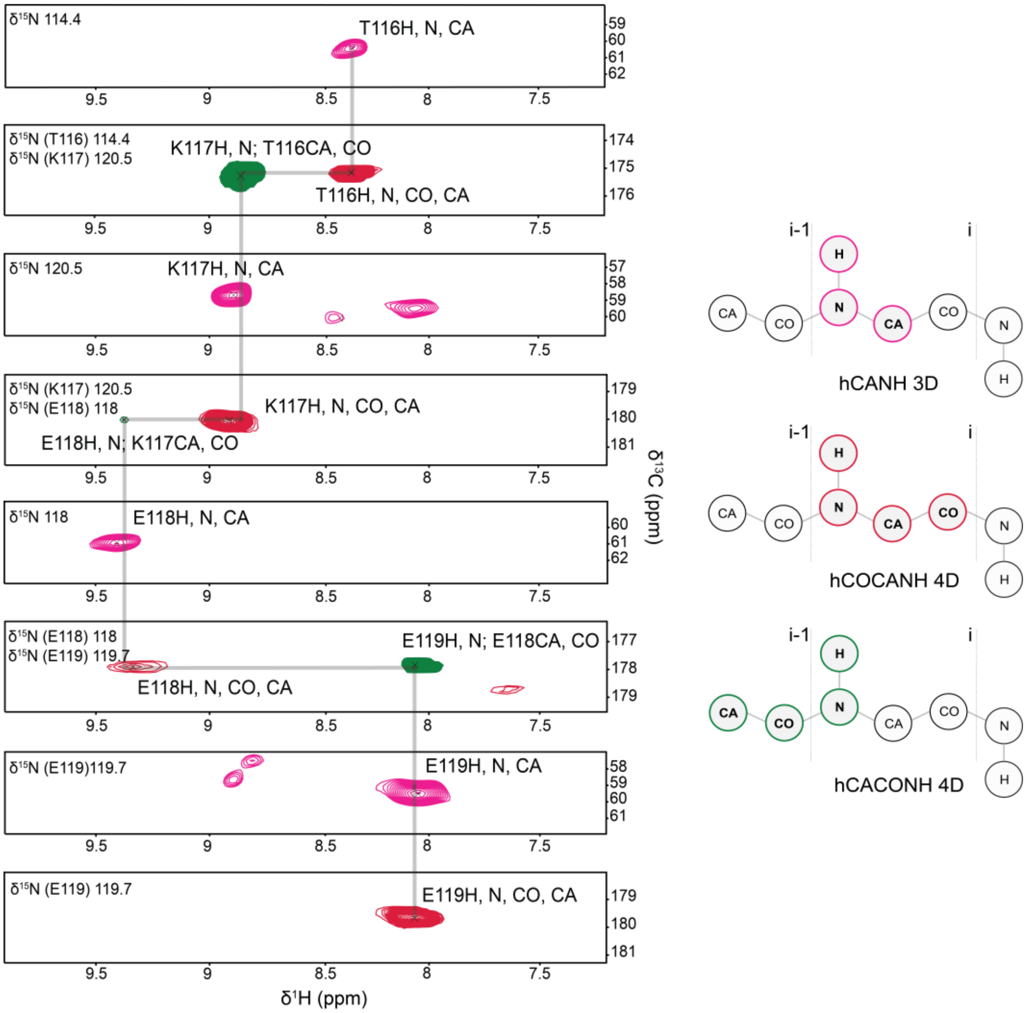


**Fig. S7. Sequential residue walk for the amino acid stretch T116 – E119 of *Ec*MscL WT by solid-state NMR.** Peaks from the 3D hCANH spectrum are shown in pink, from the 4D hCACONH spectrum in green, and from the 4D hCOCANH spectrum in red. Gray lines represent the connections used for assignments.

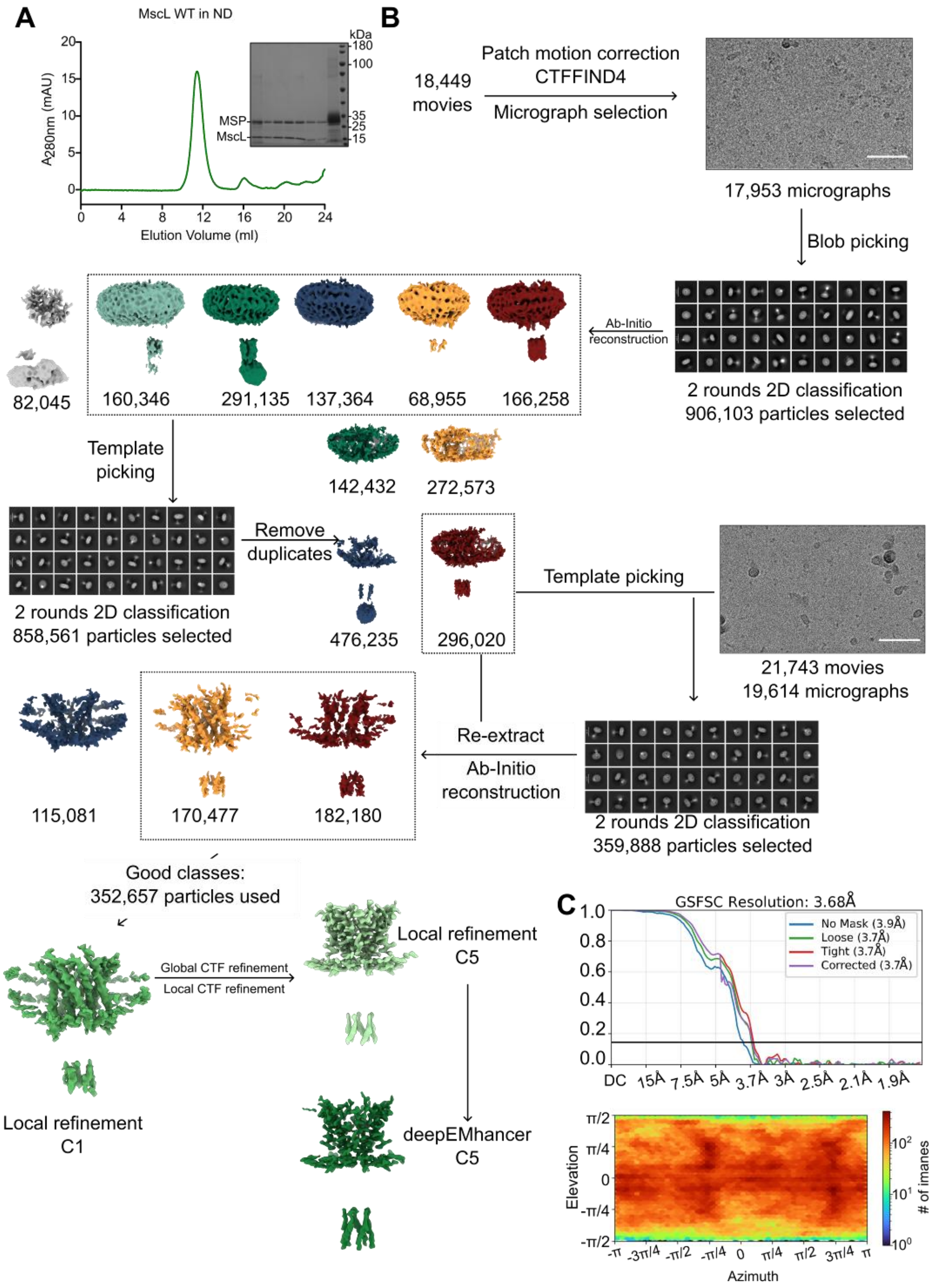


**Fig. S8. Sample preparation and Cryo-EM data processing of the *Ec*MscL wild type data using cryoSPARC v4.6.2.** (**A**) Size-exclusion chromatography and SDS-PAGE of full-length wild-type *Ec*MscL reconstituted into nanodiscs. (**B**) Workflow of image processing. Scale bar = 100 Å (**C**) The FSC curves and particle orientation distribution of the final, refined cryo-EM map of *Ec*MscL at an estimated resolution of 3.7 Å.

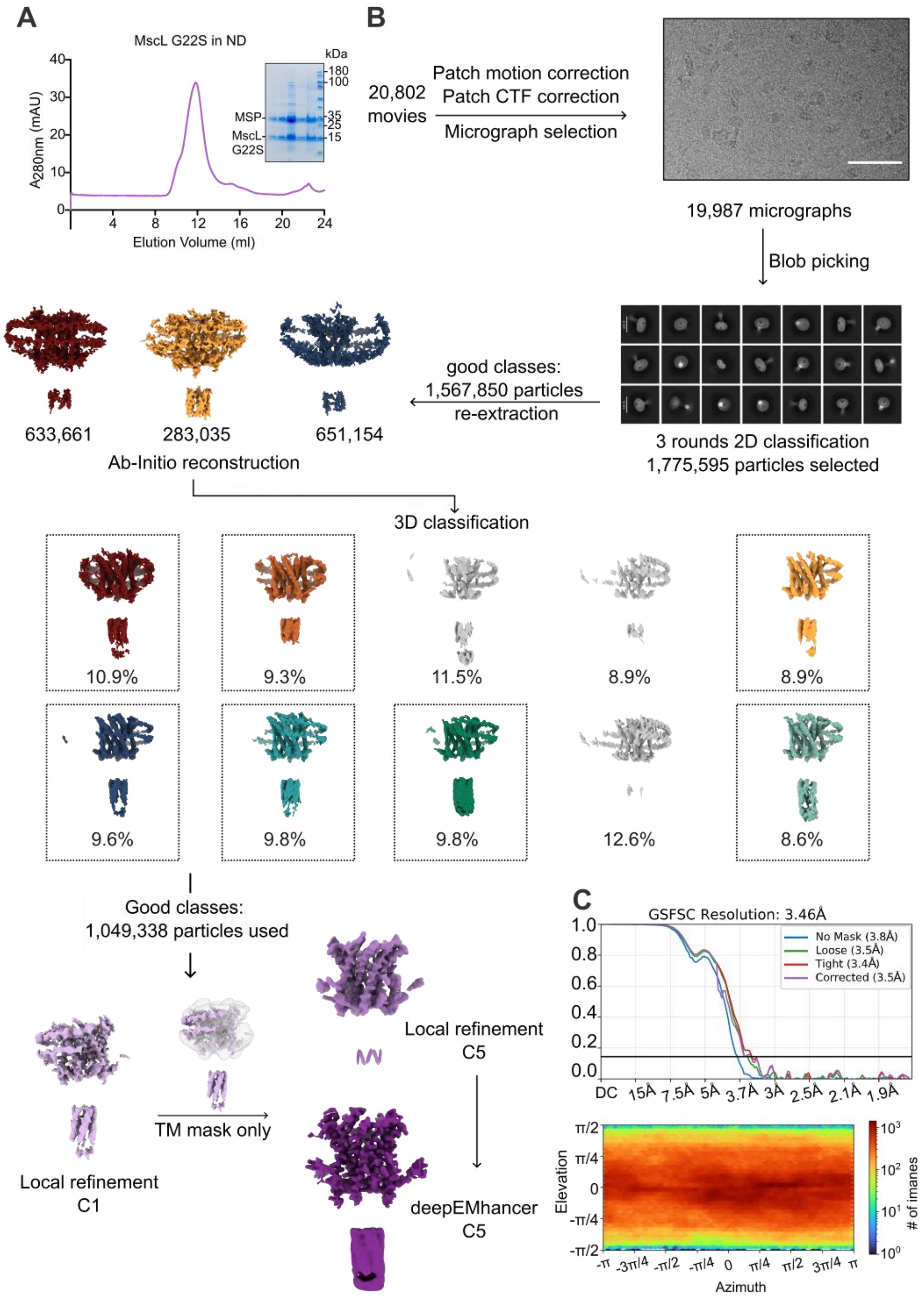


**Fig. S9. Sample preparation and data processing of the *Ec*MscL G22S mutant data using cryoSPARC v4.6.2.** (**A**) Size-exclusion chromatography and SDS-PAGE of the full-length *Ec*MscL G22S mutant reconstituted into nanodiscs. (**B**) Workflow of image processing. Scale bar = 100 Å (**C**) The FSC curves and particle orientation distribution of the final, refined cryo-EM map of *Ec*MscL G22S at an estimated resolution of 3.5 Å.

**Table S1. Assignment table for $^{13}C$,$^{15}N$-labeled WT *Ec*MscL.** Chemical shifts were referenced using internal DSS and are deposited in the BMRB (ID: 53470).

| Number | Type | CA | CB | CO | N | CD | CD1 | CG | CG1 | CG2 |
|---|---|---|---|---|---|---|---|---|---|---|
| 20 | Ala | 55.58 | | | | | | | | |
| 21 | Val | 67.1983 | 31.407 | 178.048 | 119.633 | | | | | |
| 22 | Gly | 47.8436 | | | 106.688 | | | | | |
| 23 | Val | 66.381 | | | | | | | | |
| 26 | Gly | 47.1309 | | | | | | | | |
| 27 | Ala | 54.5006 | | | | | | | | |
| 33 | Val | 67.0065 | 31.2697 | 177.264 | 119.702 | | | | | |
| 34 | Ser | 62.4876 | 62.45 | | 113.994 | | | | | |
| 35 | Ser | 59.5401 | | | 113.999 | | | | | |
| 37 | Val | 67.0382 | | | | | | | | |
| 38 | Ala | 55.4019 | | | | | | | | |
| 43 | Pro | 67.0022 | 29.637 | | | 50.2171 | | 27.4136 | | |
| 44 | Pro | 65.7797 | 30.7698 | | 132.51 | 51.1705 | | 28.402 | | |
| 45 | Leu | 58.335 | | | | | | | | |
| 46 | Gly | 47.245 | | | | | | | | |
| 57 | Phe | 58.1965 | 38.286 | 174.238 | | | | | | |
| 58 | Ala | 51.7073 | 23.032 | 175.437 | 129.468 | | | | | |
| 59 | Val | 59.7036 | 35.6563 | 173.737 | 116.608 | | | | 22.0033 | |
| 60 | Thr | 63.4583 | 68.2939 | 173.663 | 127.08 | | | | | 22.2065 |
| 61 | Leu | 55.3151 | 42.7415 | 175.81 | 126.987 | | 26.2357 | 27.2037 | | 22.851 |
| 62 | Arg | 56.1113 | 34.0984 | 174.404 | 116.895 | | | | | |
| 63 | Asp | 54.6686 | 41.2893 | 177.049 | 126.936 | 179.526 | | | | |
| 64 | Ala | 53.038 | 19.3187 | 177.4 | 124.049 | | | | | |
| 65 | Gln | 54.1328 | 30.6058 | | 121.31 | | | 33.0882 | | |
| 66 | Gly | 47.0853 | | 175.185 | | | | | | |
| 67 | Asp | 54.538 | 40.7796 | 175.822 | 127.202 | 180.525 | | | | |
| 68 | Ile | 58.609 | 37.6049 | 174.452 | 124.97 | | 12.2597 | | 27.3276 | 16.8513 |
| 69 | Pro | 62.1598 | 32.9142 | 177.291 | 138.285 | 51.2099 | | 27.2173 | | |
| 70 | Ala | 52.2619 | 19.5317 | 177.495 | 122.695 | | | | | |
| 71 | Val | 62.4625 | 32.526 | 174.953 | 123.255 | | | | 22.3023 | 20.5107 |
| 72 | Val | 60.6888 | 35.7539 | 174.037 | 129.213 | | | | 20.9478 | |
| 73 | Met | 54.2939 | 35.1237 | | 123.974 | | | | | |
| 74 | His | 56.0995 | | | | | | | | |
| 75 | Tyr | 59.842 | 38.318 | 176.79 | 118.154 | | | | | |
| 76 | Gly | 47.686 | | 175.663 | 111.607 | | | | | |
| 77 | Val | 66.6056 | 31.7693 | 177.716 | 122.383 | | | | 23.074 | 21.702 |
| 78 | Phe | 61.8022 | 38.286 | 176.837 | 119.275 | | | | | |
| 79 | Ile | 65.161 | 37.1823 | 177.338 | 117.577 | | 14.484 | | 30.5938 | 17.644 |
| 80 | Gln | 59.0604 | | 177.513 | 119.285 | | | | | |
| 81 | Asn | 56.1103 | 37.503 | | 116.349 | | | | | |
| 82 | Val | 67.136 | | | | | | | | |
| 113 | Pro | 62.6865 | | 176.373 | | | | 27.4073 | | |
| 114 | Ala | 50.0559 | 17.888 | 175.498 | 126.186 | | | | | |
| 115 | Pro | 62.5781 | 32.1869 | 177.544 | 136.232 | 50.3593 | | 27.4073 | | |
| 116 | Thr | 60.8941 | 70.6838 | 175.293 | 114.439 | | | | | 22.255 |
| 117 | Lys | 59.1759 | 31.6332 | 180.154 | 120.576 | 28.4431 | | 25.0677 | | |
| 118 | Glu | 61.4975 | 28.2815 | 177.699 | 118.4 | | | 37.355 | | |
| 119 | Glu | 60.0016 | 29.4967 | 179.788 | 120.874 | | | | | |
| 120 | Val | 67.1171 | 31.781 | 179.343 | 122.431 | | | | 22.772 | 21.197 |
| 121 | Leu | 58.2372 | 43.5005 | 179.221 | 121.421 | | | | | |
| 122 | Leu | 57.9632 | 43.528 | 178.649 | 117.98 | | | | | |
| 123 | Thr | 67.4779 | 68.5458 | 174.853 | 117.008 | | | | | 21.1589 |
| 124 | Glu | 59.9993 | 29.8734 | 179.703 | 121.881 | 183.235 | | 32.139 | | |
| 125 | Ile | 66.2729 | 38.2351 | 176.34 | 120.147 | | 14.484 | | 30.5938 | 15.7079 |
| 126 | Arg | 60.533 | 27.555 | | 120.402 | | | | | |
| 127 | Asp | 57.8565 | | | 119.839 | | | | | |

**Table S2. Assignment table for $^2H$,$^{13}C$,$^{15}N$ labelled *Ec*MscL WT.** Chemical shifts were referenced using internal DSS and are deposited in the BMRB (ID: 53468)

| Number | Type | H | N | CO | Cα |
|---|---|---|---|---|---|
| 34 | SER | 8.29 | 113.82 | 177.09 | 62.06 |
| 35 | SER | 7.59 | 114.96 | | 59.33 |
| 59 | VAL | 8.32 | 117.01 | 175.25 | 59.37 |
| 60 | THR | 9.39 | 126.66 | 173.96 | 63.18 |
| 61 | LEU | 9.11 | 127.05 | 173.59 | 54.97 |
| 62 | ARG | 7.54 | 117.16 | 176.32 | 55.59 |
| 63 | ASP | 8.85 | 126.30 | 174.46 | 54.26 |
| 64 | ALA | 8.84 | 123.54 | 176.78 | 52.77 |
| 65 | GLN | 8.32 | 121.27 | 177.38 | 53.90 |
| 66 | GLY | 9.05 | 116.40 | 175.96 | 46.99 |
| 67 | ASP | 8.98 | 126.96 | 175.19 | 54.26 |
| 68 | ILE | 8.28 | 124.77 | 175.79 | 58.07 |
| 70 | ALA | 8.72 | 121.70 | 177.01 | 51.99 |
| 71 | VAL | 8.89 | 122.39 | 177.94 | 62.13 |
| 72 | VAL | 8.88 | 128.78 | 174.77 | 60.32 |
| 73 | MET | 9.11 | 123.81 | 173.85 | 53.85 |
| 74 | HIS | 8.22 | 117.89 | 176.29 | 57.20 |
| 75 | TYR | 7.87 | 118.20 | | 59.47 |
| 76 | GLY | 7.80 | 111.33 | | 47.24 |
| 77 | VAL | 7.51 | 121.99 | 175.41 | 65.97 |
| 79 | ILE | 8.40 | 117.84 | 177.09 | 64.75 |
| 80 | GLN | 8.16 | 119.37 | 178.59 | 58.88 |
| 81 | ASN | 7.96 | 116.34 | 177.31 | 56.19 |
| 113 | PRO | | | 176.36 | |
| 114 | ALA | 9.09 | 126.132 | 174.36 | 50.52 |
| 115 | PRO | | | 177.70 | 62.79 |
| 116 | THR | 8.35 | 114.27 | 177.72 | 60.40 |
| 117 | LYS | 8.92 | 120.36 | 175.22 | 58.69 |
| 118 | GLU | 9.44 | 118.10 | 180.05 | 60.93 |
| 119 | GLU | 8.09 | 119.87 | 177.76 | 59.50 |
| 120 | VAL | 8.36 | 122.35 | 179.82 | 66.60 |
| 121 | LEU | 8.02 | 121.19 | 179.34 | 57.92 |
| 122 | LEU | 8.83 | 117.71 | 179.35 | 57.53 |
| 123 | THR | 8.17 | 116.77 | 178.70 | 67.09 |
| 124 | GLU | 7.87 | 121.29 | 174.91 | 59.59 |
| 125 | ILE | 8.45 | 119.3 | 176.33 | 66.2 |
| 126 | ARG | 8.44 | 120.22 | 178.71 | 60.10 |
| 127 | ASP | 8.84 | 119.83 | 178.97 | 57.50 |
| 128 | LEU | 8.76 | 121.34 | 178.98 | 59.27 |

**Table S3. Assignment table for $^{2}$H,$^{13}$C,$^{15}$N labelled *Ec*MscL G22S.** Chemical shifts were referenced using internal DSS and are deposited in the BMRB (ID: 53469)

| Number | Type | H | N | CO | Cα | Cβ |
|---|---|---|---|---|---|---|
| 33 | VAL | 8.31 | 119.05 | 177.05 | 66.94 | |
| 34 | SER | 8.22 | 113.82 | | 61.76 | |
| 58 | ALA | 8.51 | 130.11 | | 51.58 | |
| 62 | ARG | 8.38 | 117.40 | 177.25 | 55.70 | |
| 63 | ASP | 9.14 | 125.78 | 174.35 | 53.55 | |
| 64 | ALA | 8.92 | 122.00 | | 54.17 | |
| 73 | MET | 9.15 | 124.51 | | 54.01 | |
| 77 | VAL | 8.15 | 121.61 | 179.31 | 66.16 | |
| 78 | PHE | 8.27 | 119.02 | 178.88 | 61.24 | 42.31 |
| 79 | ILE | 8.41 | 118.03 | 177.46 | 64.074 | |
| 80 | GLN | 8.08 | 119.22 | 178.47 | 59.29 | 27.66 |
| 81 | ASN | 8.55 | 116.45 | | 56.57 | |
| 113 | PRO | | | 176.36 | | |
| 114 | ALA | 8.48 | 125.66 | | 49.97 | 25.30 |
| 116 | THR | 8.41 | 113.85 | 177.63 | 60.40 | |
| 117 | LYS | 8.94 | 120.79 | 175.17 | 58.82 | |
| 118 | GLU | 9.18 | 117.63 | 179.67 | 60.57 | |
| 119 | GLU | 8.18 | 120.10 | | 59.16 | |
| 120 | VAL | 8.44 | 122.66 | 179.74 | 66.61 | |
| 121 | LEU | 8.14 | 121.04 | 179.34 | 57.71 | 42.31 |
| 122 | LEU | 8.74 | 117.52 | 179.46 | 57.41 | 42.32 |
| 123 | THR | 8.17 | 117.12 | 178.56 | 67.12 | |
| 124 | GLU | 7.89 | 121.26 | 175.02 | 59.42 | |
| 125 | ILE | 8.57 | 119.85 | 179.59 | 65.80 | |
| 126 | ARG | 8.46 | 120.14 | | 60.02 | |
| 127 | ASP | 8.82 | 119.93 | 178.89 | 57.47 | 38.73 |
| 128 | LEU | 8.73 | 121.21 | 179.06 | 59.49 | |

**Table S4. Statistics for data collection and refinement.** Statistics for 3D reconstruction (cryoSPARC 4.6.2), model refinement (*Coot* 0.9.6.2 and PHENIX 1.21.2), and validation (PHENIX 1.21.2) for the cryo-EM structural refinement of *Ec*MscL wild type (WT) and G22S mutant.

| | *Ec*MscL WT | *Ec*MscL G22S |
|---|---|---|
| | PDB ID: 9TIU | PDB ID: 9TIV |
| **Data collection** | | |
| Microscope | Titan Krios G3i | Titan Krios G3i |
| Detector | Gatan K3 | Gatan K3 |
| Energy filter slit width (eV) | 20 | 20 |
| Magnification | 105,000 | 105,000 |
| Voltage (kV) | 300 | 300 |
| Electron exposure ($e^{-}/Å^{2}$) | 60 | 60 |
| Defocus range (μm) | -2.8~-1 | -2.4~-0.8 |
| Pixel size (Å) | 0.83 | 0.83 |
| Symmetry imposed | C5 | C5 |
| Number of micrographs | 37,567 | 19,987 |
| Initial particle images (no.) | 22,160,563 | 9,266,266 |
| Final particle images (no.) | 352,657 | 1,049,338 |
| Map resolution (Å) | | |
| FSC 0.5(unmasked/masked) | 6.5/4.1 | 4.1/4.1 |
| FSC 0.143(unmasked/masked) | 3.7/3.6 | 3.5/3.4 |
| Sharpening B-factor ($Å^{2}$) | −116.2 | −182.0 |
| **Model refinement** | | |
| Number of atoms | | |
| All (hydrogens) | 8355(4175) | 9190(4595) |
| Protein residues | 575 | 630 |
| Ligand (sugars) | 0 | 0 |
| Model validation | | |
| CC (mask) map versus model (%) | 0.73 | 0.77 |
| CC (box) map versus model (%) | 0.62 | 0.65 |
| RMSD | | |
| Bond lengths (Å) | 0.003 | 0.002 |
| Bond angles (°) | 0.598 | 0.568 |
| Ramachandran statistics | | |
| Favored regions (%) | 98.35 | 98.03 |
| Allowed regions (%) | 1.65 | 1.97 |
| Outliers (%) | 0 | 0 |
| Rotamer outliers (%) | 6.41 | 1.38 |
| Cβ outliers (%) | 0.93 | NA |
| CaBLAM outliers (%) | 0.00 | 1.69 |
| Clashscore | 10.31 | 8.60 |
| MolProbity overall score | 2.15 | 1.57 |

**Table S5. Parameters for all magnetization transfer steps.** Sensitivity-enhanced 3D and 4D experiments used for assignments of *Ec*MscL WT. The nutation frequency is given for the maximum amplitude of each pulse. Experiments were conducted at 900 MHz.

| Experiment | hCANH | hCONH | hCACONH | hCOCANH |
|---|---|---|---|---|
| **$^{1}H$-$^{13}C$** | | | | |
| $^{1}H$ shape | ramp 90-100 | ramp 90-100 | ramp 90-100 | ramp 90-100 |
| $^{1}H$ power | 82 kHz (8.2 W) | 83 kHz (8.3 W) | 82 kHz (8.2 W) | 83 kHz (8.3W) |
| $^{13}C$ shape | rectangular | rectangular | rectangular | rectangular |
| $^{13}C$ power | 20 kHz (1.8 W) | 20 kHz (1.8 W) | 20 kHz (1.8 W) | 20 kHz (1.8 W) |
| duration (ms) | 2.7 | 2.8 | 2.8 | 2.8 |
| **$^{13}C$-$^{13}C$ (backbone)** | | | | |
| $^{13}C$ shape | - | - | homo TROP | homo TROP |
| $^{13}C$ power | - | - | 68 kHz (21 W) | 73 kHz (24 W) |
| duration (ms) | - | - | 1.65 | 1.65 |
| **$^{13}C$-$^{15}N$** | | | | |
| $^{13}C$ shape | TROP | TROP | TROP | TROP |
| $^{13}C$ power | 69 kHz (22 W) | 69 kHz (22 W) | 69 kHz (22 W) | 69 kHz (22 W) |
| $^{15}N$ shape | TROP | TROP | TROP | TROP |
| $^{15}N$ power | 46 kHz (50 W) | 46 kHz (50 W) | 46 kHz (50 W) | 46 kHz (50 W) |
| duration (ms) | 3.33 | 3.33 | 3.33 | 3.33 |
| **$^{15}N$-$^{1}H$** | | | | |
| $^{15}N$ shape | rectangular | rectangular | TROP | TROP |
| $^{15}N$ power | 14 kHz (4.5 W) | 14 kHz (4.5 W) | 52.6 kHz (65 W) | 52.6 kHz (65 W) |
| $^{1}H$ shape | ramp 80-100 | ramp 80-100 | TROP | TROP |
| $^{1}H$ power | 75 kHz (8 W) | 75 kHz (8 W) | 52 kHz (3.9 W) | 52 kHz (3.9 W) |
| duration (ms) | 0.7 | 0.7 | 0.7 | 0.7 |

**Table S6. Parameters for all magnetization transfer steps.** Sensitivity-enhanced 3D and 4D experiments used for assignments of *Ec*MscL G22S. The nutation frequency is given for the maximum amplitude of each pulse. Experiments were conducted at 600 MHz, unless stated otherwise.

| Experiment | hCANH | hCONH | hCAcoNH | hCOcaNH | hCXCANH (900 MHz) |
|---|---|---|---|---|---|
| **$^{1}$H-$^{13}$C** | | | | | |
| $^{1}$H shape | ramp 90-100 | ramp 90-100 | ramp 90-100 | ramp 90-100 | ramp 90-100 |
| $^{1}$H power | 79 kHz (3.6 W) | 79 kHz (3.6 W) | 80 kHz (3.7 W) | 81 kHz (3.8 W) | 80 kHz (9.2 W) |
| $^{13}$C shape | rectangular | rectangular | rectangular | rectangular | rectangular |
| $^{13}$C power | 20 kHz (1.8 W) | 20 kHz (1.8 W) | 20 kHz (1.8 W) | 20 kHz (1.8 W) | 20 kHz (1.8 W) |
| duration (ms) | 1.8 | 1.8 | 1.8 | 1.8 | 6 |
| **$^{13}$C-$^{13}$C (side-chains)** | | | | | |
| $^{13}$C shape | | | | | DIPSI 3 |
| $^{13}$C power | | | | | 15.62 kHz (1.12 W) |
| duration (ms) | | | | | 0.016 |
| **$^{13}$C-$^{13}$C (backbone)** | | | | | |
| $^{13}$C shape | - | - | homo TROP | homo TROP | |
| $^{13}$C power | - | - | 73 kHz (24 W) | 73 kHz (24 W) | |
| duration (ms) | - | - | 1.8 | 1.8 | |
| **$^{13}$C-$^{15}$N** | | | | | |
| $^{13}$C shape | TROP | TROP | TROP | TROP | TROP |
| $^{13}$C power | 67 kHz (20.2 W) | 66 kHz (19.5 W) | 66 kHz (19.5 W) | 67 kHz (20.2 W) | 70 kHz (22 W) |
| $^{15}$N shape | TROP | TROP | TROP | TROP | TROP |
| $^{15}$N power | 42 kHz (21.3 W) | 42 kHz (21.3 W) | 42 kHz (21.3 W) | 42 kHz (21.3 W) | 46 kHz (50 W) |
| duration (ms) | 3.64 | 3.64 | 3.64 | 3.64 | 3.33 |
| **$^{15}$N-$^{1}$H** | | | | | |
| $^{15}$N shape | TROP | TROP | TROP | TROP | TROP |
| $^{15}$N power | 55 kHz (36 W) | 55 kHz (36 W) | 55 kHz (36 W) | 55 kHz (36 W) | 52.6 kHz (65 W) |
| $^{1}$H shape | TROP | TROP | TROP | TROP | TROP |
| $^{1}$H power | 61 kHz (2.1 W) | 61 kHz (2.1 W) | 61 kHz (2.1 W) | 61 kHz (2.1 W) | 52 kHz (3.9 W) |
| duration (ms) | 0.8 | 0.8 | 0.8 | 0.8 | 0.733 |

**Table S7. Acquisition parameters.** Sensitivity-enhanced 3D and 4D experiments used for the assignments of *Ec*MscL WT. SW = spectral width, NUS = Non-uniform sampling.

| Experiment | hCANH | hCONH | hCACONH | hCOCANH |
|---|---|---|---|---|
| Scans | 56 | 80 | 80 | 112 |
| $^{1}$H acquisition time (ms) | 30.7 | 30.7 | 24 | 24 |
| $^{1}$H points | 3072 | 3072 | 2400 | 2400 |
| $^{1}$H SW (ppm) | 55.5 | 55.5 | 55.5 | 55.5 |
| $^{13}$CA acquisition time (ms) | 4.2 | - | 4.6 | 4.6 |
| $^{13}$CA points | 54 | - | 48 | 48 |
| $^{13}$CA SW (ppm) | 28 | - | 23 | 23 |
| $^{13}$CO acquisition time (ms) | - | 5.6 | 6.6 | 6.6 |
| $^{13}$CO points | - | 36 | 36 | 36 |
| $^{13}$CO SW (ppm) | - | 14 | 12 | 12 |
| $^{15}$N acquisition time (ms) | 11.3 | 7.8 | 9.8 | 9.8 |
| $^{15}$N points | 58 | 40 | 36 | 36 |
| $^{15}$N SW (ppm) | 28 | 28 | 20 | 20 |
| NUS points | - | - | 387 | 386 |
| NUS % | - | - | 5 | 5 |
| total points | 3132 | 1440 | 3096/62208 | 3088/62208 |
| Experimental time (h) | 65 h | 43 h | 140 h | 196 h |

**Table S8. Acquisition parameters.** Sensitivity-enhanced 3D and 4D experiments used for the assignments of *Ec*MscL G22S. SW = spectral width, NUS = Non-uniform sampling.

| Experiment | hCANH | hCONH | hCAcoNH | hCOcaNH | hCXCANH (900 MHz) |
|---|---|---|---|---|---|
| Scans | 152 | 128 | 192 | 304 | 120 |
| $^{1}$H acquisition time (ms) | 30 | 30 | 30 | 30 | 18.8 |
| $^{1}$H points | 1438 | 1438 | 1438 | 1438 | 1344 |
| $^{1}$H SW (ppm) | 40 | 40 | 40 | 40 | 39.7 |
| $^{13}$CA acquisition time (ms) | 5.3 | - | 4.6 | - | 4 |
| $^{13}$CA points | 48 | - | 42 | - | 40 |
| $^{13}$C SW (ppm) | 30 | - | 30 | - | 22 |
| $^{13}$CO acquisition time (ms) | - | 8.8 | - | 8.8 | - |
| $^{13}$CO points | - | 32 | - | 32 | - |
| $^{13}$CO SW (ppm) | - | 12 | - | 12 | - |
| $^{13}$CX acquisition time (ms) | - | - | - | - | 3.9 |
| $^{13}$CX points | - | - | - | - | 116 |
| $^{13}$CX SW (ppm) | - | - | - | - | 65 |
| $^{15}$N acquisition time (ms) | 14.6 | 12.7 | 12.7 | 12.7 | 9.2 |
| $^{15}$N points | 48 | 42 | 42 | 42 | 32 |
| $^{15}$N SW (ppm) | 27 | 27 | 27 | 27 | 19 |
| NUS points | - | - | - | - | 739 |
| NUS % | - | - | - | - | 4 |
| total points | 2304 | 1344 | 1764 | 1344 | 5 912/148480 |
| Experimental time (h) | 136 h | 66 | 153 | 153 | 265 h |
